# Insights into calcium signaling and gene expression in astrocytes uncovered with 129S4 Slc1a3-2A-CreERT2 knock-in mice

**DOI:** 10.1101/2020.04.06.027714

**Authors:** Lech Kaczmarczyk, Nicole Reichenbach, Nelli Blank, Maria Jonson, Lars Dittrich, Gabor C. Petzold, Walker Jackson

## Abstract

Genetic variation is a primary determinant of phenotypic diversity within populations. In laboratory mice, genetic variation has often been regarded as a serious experimental confounder, and thus minimized through inbreeding. However, generalizations of results obtained with inbred strains need to be made with caution. Effects of genetic background on traits need to be controlled, especially when working with complex phenotypes and disease models. Here we compared behavioral parameters of C57Bl/6 – the mouse strain most widely used for biomedical research - with those of 129S4. Our data demonstrate high within-strain and intra-litter behavioral hyperactivity in C57Bl/6. In contrast, 129S4 had relatively consistent activity levels throughout life. This consistency would be advantageous for studying neurodegeneration and aging, when mice need to be analyzed for long periods. However, the majority of mouse models and transgenic tools are on a C57Bl/6 background. We recently established six popular Cre driver lines and two Cre effector lines in 129S4. To augment this collection, we genetically engineered a Cre mouse line to study astrocytes directly in 129S4, which we describe here. For functional validation, it was crossed with two Cre effector lines, each in a different genomic locus, and showed in both cases that it was functional and astrocyte-specific. Calcium currents studied with gCaMP5g-tdTomato were more heterogenous, lasted longer and had a higher amplitude in cortical compared to hippocampal astrocytes. Translatomes studied with RiboTag revealed that some genes thought to mark neurons are also expressed in astrocytes, that genes linked to a single neurodegenerative disease have highly divergent expression patterns, and that ribosome proteins are non-uniformly expressed across brain regions and cell types.

## Introduction

Humans are highly diversified on multiple levels. The high variability between individuals and groups influences vulnerability to diseases and responsiveness to medical treatments. Phenotypic differences are reflected in gene- and protein expression profiles and similar variations are observed in other mammals, including laboratory mice (Wade and Daly, 2005). Genetic background is a primary factor affecting disease phenotypes in mouse models (Erickson, 1996; Lloret et al., 2006; Marques et al., 2011) and a notorious experimental confounder. Therefore, inbred mouse strains, in which all individuals are genetically identical, are often employed to limit phenotypic variation. C57Bl/6 is the most commonly used inbred mouse strain with over 44000 publications on Pubmed as of February 2020 (Bryant, 2011; Sarsani et al., 2019). C57Bl/6 includes numerous sub-strains, collectively known as Black 6 or B6. Their popularity stems from some important features. First, since they carry a recessive mutation resulting in black fur, they were widely used during the early days of gene-targeting experiments as sources of host blastocysts and breeding partners for chimeras. This provided an invaluable screening tool for quick exclusion of non-chimeric mice and progeny of chimeric mice that did not inherit the desired genetically engineered ES cell genome, simply by inspecting fur colors, a convenience still widely used today. Second, B6 mice are more susceptible than most other strains to certain drugs and self-administer them when given the chance (Elmer et al., 2010; Orsini et al., 2005; Yoneyama et al., 2008). They also perform well in learning and memory paradigms, as they are inclined to explore more than other strains (Lhotellier et al., 1993; Sultana et al., 2019; Tarantino et al., 2000). Finally, B6 mice are very active, which is a useful characteristic for studying metabolic rates and effects of exercise on the body (Funkat et al., 2004).

Congruent with these characteristics, we observed hyperactivity in a subset of mice congenic for a B6 background (C57Bl/6NTac), especially at night (Jackson et al., 2009; Jackson et al., 2013). Since the hyperactivity was observed in knock-in controls, we suspected it was due to the genetic background rather than the engineered mutations. This was alarming since nocturnal hyperactivity in only a subset of mice would result in those individuals being more tired than the remaining population during the day, which would likely go unnoticed although affecting sleep state and gene expression. We also observed an obese subset which was troublesome since obesity also leads to gene expression differences in the brain (Siino et al., 2018). A subset of individuals of an inbred strain with atypical characteristics makes the population heterogeneous, diminishing the value of using an inbred strain. We therefore tested an alternative genetic background, 129S4 (formerly 129Sv/Jae, hereafter S4). This strain is used in amphetamine research due to its calm nature (Crittenden et al., 2014; Crittenden et al., 2019). Moreover, a quantitative electroencephalography (EEG) study revealed several differences in sleep physiology between S4 and B6 (Dittrich et al., 2017). Aberrations in sleep phenotypes are linked to diseases, including neurodegeneration, cancer, and diabetes (Aalling et al., 2018; Chen et al., 2018; Hakim et al., 2014; Malhotra, 2018), indicating that sleep profoundly affects pathology. Consequently, the differences between mouse strains in this domain might translate to strain-dependent experimental differences in mouse models of disease. This has two important implications. First, it highlights the importance of choosing the optimal genetic background for studying specific phenotypic parameters. Second, it suggests that using mouse strains with disparate characteristics (e.g., B6 vs S4 with respect to sleep phenotypes) can be informative in identifying gene expression mechanisms that function independent of genetic background. Such an analysis would justify generalizing experimental conclusions beyond individual strains.

Here, using automated video tracking analysis, we systematically compared the home cage behavior of S4 and B6 mice and verified the pronounced intra-strain and within-litter variabilities in B6. We expected these differences to affect phenotypes in mouse disease models, and to mask subtle gene expression characteristics of pre-clinical phases of neurodegenerative and psychiatric disorders. Therefore, in recent years we have backcrossed multiple Cre and Cre reporter lines to S4 (Kaczmarczyk et al., 2019), but that was very time consuming. Unfortunately, an astrocyte-specific Cre line we previously backcrossed to S4 to study stroke (Rakers et al., 2019) was impractical to use because mice could not be maintained as homozygotes (Kretz et al., 2003). To meet the need for a better Cre line for astrocytes while avoiding extensive backcrossing of existing lines we generated an isogenic 129S4 inducible Cre driver line. These mice express CreERT2 (Feil et al., 2009) from the solute carrier family 1 member 3 (Slc1a3) locus (Shibata et al., 1997) while preserving expression of the endogenous protein using the 2A peptide bicistronic expression method (Szymczak et al., 2004). The ability of Slc1a3 gene elements to drive high and specific transgene expression in astrocytes has been previously demonstrated (Anthony and Heintz, 2008; Mori et al., 2006; Regan et al., 2007; Slezak et al., 2007), but a knock-in approach to express a transgene from the endogenous *Slc1a3* locus while preserving expression of the endogenous protein has not been reported. Here we attempted to fill this void. We demonstrate the functionality of the Slc1a3-2A-CreERT2 mouse line with two separate Cre reporter lines. The GCaMPg5 calcium sensor revealed functional differences between hippocampal and cortical astrocytes *in vivo*. RiboTag mice were used to compare translating mRNAs from cerebellar astrocytes, almost exclusively comprising Bergmann glia, with those of the rest of the brain, providing insights into the translatomes of astroglia in these different regions.

## Results

### A subset of B6 mice are hyperactive at night

We performed video-based behavior recognition experiments to determine behavioral characteristics of B6 and S4 strains. Mice were video recorded in standard mouse cages for 24 hr periods every 2 months starting at 6 months of age. The videos were scored automatically by computer software as described previously (Jackson et al., 2009; Steele et al., 2007). A comparison across ages revealed many differences in activities of adult S4 and B6 mice, even into old age (Fig 1A). Importantly, a closer inspection of data from individual mice revealed a subset of B6 mice demonstrating hyper-repetitive activities for behaviors requiring exertion (e.g., jumping, walking, rearing, hanging, etc.) and also feeding (Fig 1B). Moreover, individuals hyperactive in one behavior were often not hyperactive in others. For example, in 24 hours, one mouse traveled 4,000 meters whereas another mouse jumped for over 3 hours and yet another mouse kept its snout in the food bin “eating” for 3.8 hours (Fig S1). Interestingly, another subset of B6 became much fatter than the rest (Fig 1B). For example, at 18 months 5 of 25 B6 mice were over 20% heavier than the median weight (42.4 g), 4 of which were over 30% heavier. The fat B6 mice were not the same as those with the highest eating scores (Fig S1), indicating the high eating scores included activities besides consumption of food, possibly excessive gnawing. Remarkably, much of the hyper-repetitive behaviors occurred during the dark cycle, when mice are typically not observed by human investigators (Movie S1, https://tinyurl.com/wcmz9zf). In contrast, S4 mice demonstrated no hyper-repetitive behaviors and 0 of 17 S4 mice became more than 20% heavier than the median weight (29.9 g) despite eating the same standard chow as the B6 mice. Such drastically different behaviors and weights likely have an impact on metabolism and physiology, and finally on gene expression (Nadler et al., 2006). Therefore, to reduce the inter-replicate variability in gene expression studies, we have begun to use the S4 genetic background in most of our experiments.

**Figure 1.**
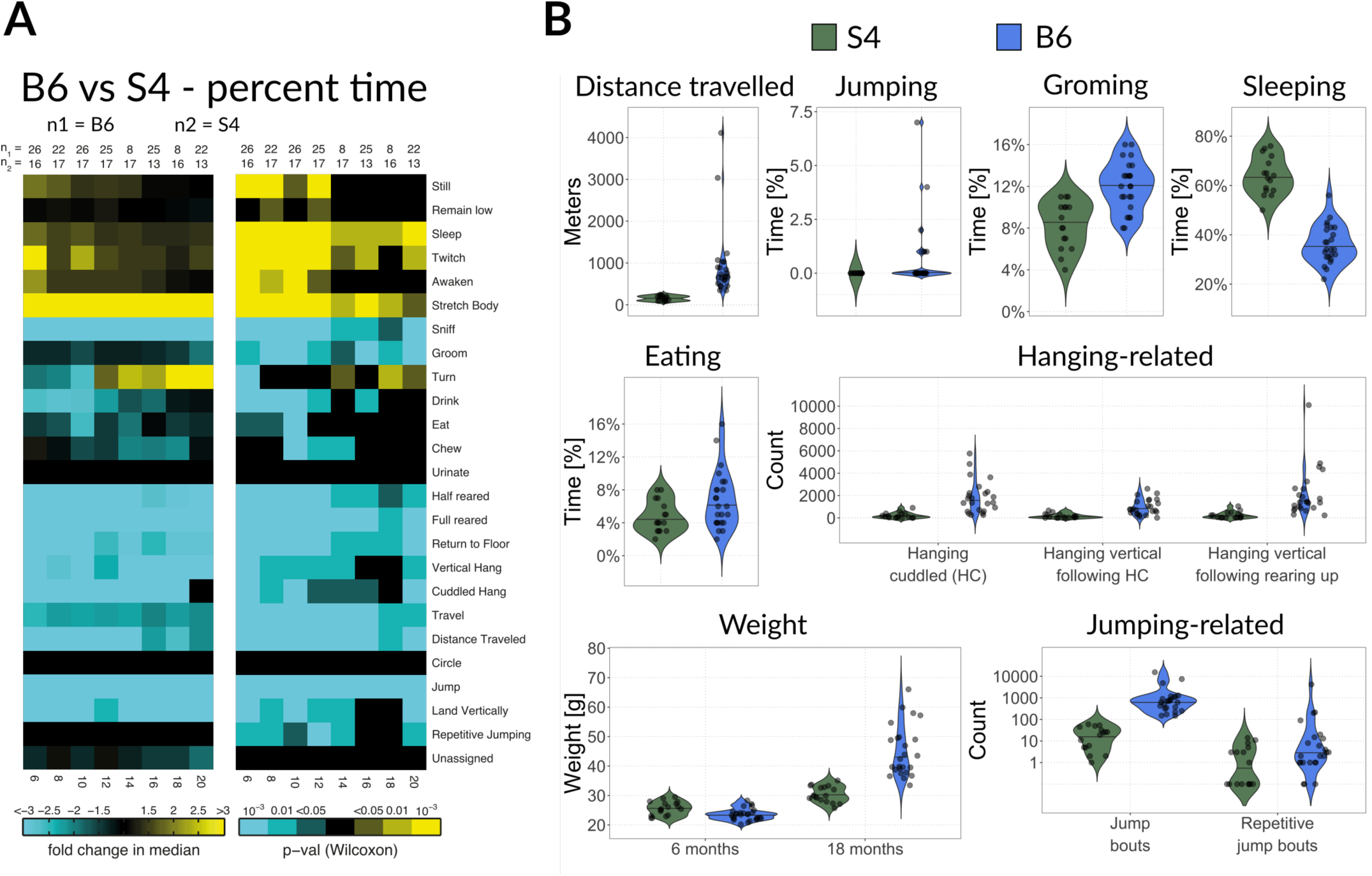
Home cage automated behavioral analysis. (**A**) Heat map showing statistical summary of longitudinal video monitoring analysis performed at 6, 8, 10, 12, 14, 16, 18, and 20 months of age on female mice. (**B**) Violin plot representation of the behavioral data from A at 6 months. For weight comparison, a cohort measured at 18-month time point was additionally included. Note the scale is log_10_ for the “Jumping-related” panel

### Generation of the S4 Slc1a3-2A-CreERT2 mouse line

During brain development, neural precursor cells (NPCs) express many genes typically expressed by mature astrocytes (Ferenczy et al., 2013; Foo and Dougherty, 2013). Therefore, constitutive expression of Cre with astrocyte specific marker genes or promoter elements results in recombination in cells that later become neurons (Requardt et al., 2009; Zhang et al., 2013). To avoid this ectopic activation, we used an inducible Cre variant, CreERT2, that can be chemically activated after development to circumvent the problem of neuronal targeting (Feil et al., 2009; Slezak et al., 2007). We chose to use a knock-in approach since bacterial artificial chromosome (BAC) transgenes can be highly disruptive to the genome, both through the insertion of a very large transgene and massive deletions at the integration site (Goodwin et al., 2019; Kaczmarczyk and Jackson, 2015). Moreover, BAC transgenes often carry additional genes due to their large size. To identify a native gene to carry the CreERT2 coding sequence we evaluated the native expression patterns of three genes encoding well known astrocyte marker proteins, Aldh1l1 (Aldehyde dehydrogenase 1 family member l1), Gja1 (Gap junction alpha 1, also known as connexin 43) and Slc1a3, also known as glutamate aspartate transporter (GLAST) or excitatory amino acid transporter 1 (EAAT1), with mRNA in situ hybridization. Although Aldh1l1 was weakly detected (not shown), Gja1 and Slc1a3 demonstrated strong expression throughout the brain, very similar to that presented in the Allen Brain Atlas database (https://mouse.brain-map.org/). Although the overall expression of Gja1 was higher than that of Slc1a3, we saw two important advantages with Slc1a3. First, the cerebellar expression of Gja1 was prominent in the Purkinje cell layer (the location of Bergmann glia, BG), in the granule cell layer and in the deep cerebellar nucleus (Fig 2A). In contrast, cerebellar expression of Slc1a3 was restricted to BG (Fig 2A). We thought that having a Cre active in a very homogeneous cell type in an easily accessed brain region would be extremely useful, especially for gene expression studies. Second, the homozygous knock-out state is lethal for Gja1 but not Slc1a3, so that in the event of partial gene inactivation of Slc1a3, the health impact for homozygotes should be negligible.

**Figure 2.**
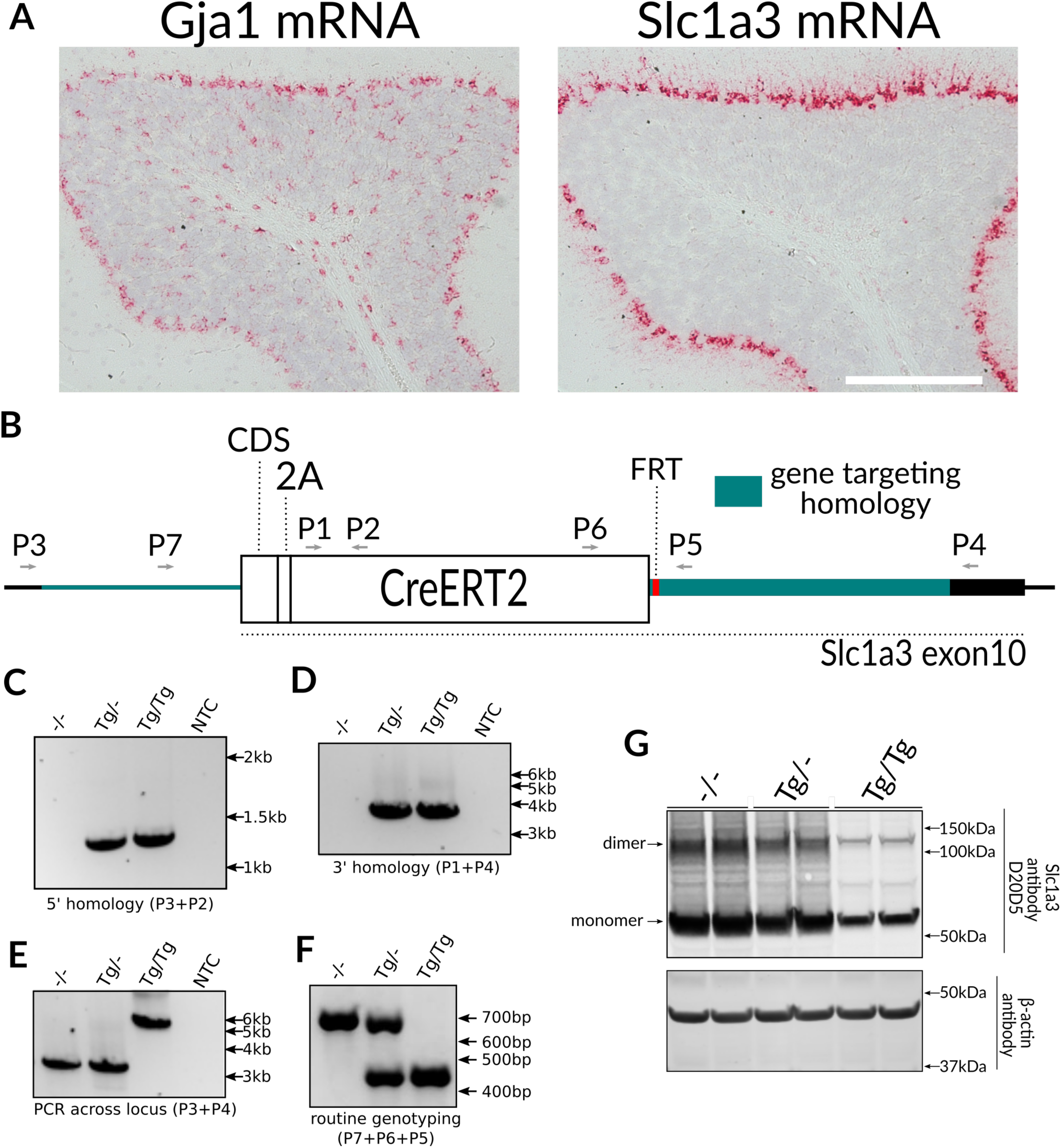
Generation of Slc1a3-2A-CreERT2 mice in 129S4 background. (**A**) In situ hybridization for Gja1 and Slc1a3. Note disparate expression pattern in the cerebellum, where Slc1a3 targets almost exclusively Bergman glia. (B) Schematic of Slc1a3-2A-CreERT2 targeted locus (elements are to scale). (**C, F**) Genotyping PCR. PCRs with primers flanking 5; homology region (**C**), 3’ homology region (**D**), flanking the whole targeted locus (we never managed to obtain the targeted band for heterozygote, probably due to WT target preference of the assay), (**E**), and routine genotyping assay (**F**) were shown. (**G**) Immunoblot showing Slc1a3 protein levels in WT, heterozygous and homologous knock-in mice.

To express CreERT2 from the *Slc1a3* locus while preserving expression of the endogenous gene product, we used the 2A peptide strategy (Fig 2B). A pair of Cas9 nickases targeted to the proximity of the STOP codon were used to ensure efficient targeting with short (1.1 kb left and 1.6 kb right) homology arms. A positive ES cell selection cassette (FRT-NeoR-FRT, FNF) was placed downstream of CreERT2 in the 3’ untranslated region (UTR). The FNF cassette was later removed by breeding to ROSA26-Flpo deleter mice on the S4 background (Raymond and Soriano, 2007). To verify gene targeting fidelity, we performed PCR across each homology region (Fig 2C and 2D), as well as across the whole locus, using primers located outside of the regions present in the targeting vector (Fig 2E). Finally, we verified that wild type Slc1a3 was absent in homozygous Slc1a3-2A-CreERT2 mice (Fig 2F). All this unambiguously verified that no undesirable duplication or O-type recombination took place (Valancius and Smithies, 1991). Immunoblotting on total brain lysates showed expression of Slc1a3 protein was markedly decreased in heterozygotes, and further decreased in homozygotes (Fig 2G). By comparing the relative counts of sequencing reads mapping to Slc1a3 and CreERT2 coding sequences in heterozygous mice, we determined that mRNA levels are only mildly affected by the presence of the transgene, indicating the reduced expression is caused by a post-transcriptional effect. Importantly, homozygous mice appear and behave normally and we maintain the line as homozygous. To test for reproductive issues, we compared the litter sizes of all S4 lines that we keep (10 in total), and observed no differences (Fig S2). In the experiments that follow, homozygous Slc1a3-2A-CreERT2 mice were crossed with Cre reporter strains such that experimental mice were hemizygous for both transgenes.

### Calcium currents are highly dynamic in cortical but not hippocampal astrocytes

The CreERT2 fusion protein is engineered to be anchored in the cytoplasm through binding with heat shock protein 90 (Hsp90) (Feil et al., 2009). In the presence of 4-hydroxytamoxifen (4-OHT) Hsp90 releases CreERT2 which is then promptly shuttled into the nucleus for recombination. Peripheral application of tamoxifen also works as it is processed to 4-OHT by the liver. To test the functionality of our Slc1a3-2A-CreERT2 line we bred them to a Cre reporter line, PC::G5-tdT, that encodes a green fluorescent protein fused to the calcium binding protein calmodulin (GCaMP5), useful for measuring changes to intracellular calcium levels (Gee et al., 2014). This transgene also encodes a separate red fluorescent protein, tdTomato, that provides stable fluorescence, independent of calcium fluctuations, thereby serving as a baseline reference for fluorescence emission. We activated expression of this construct (G5-tdT) in three months old mice with tamoxifen.

We first determined if activation was specific to astrocytes and to what extent astrocytes were expressing the reporter. Therefore, we labeled formaldehyde fixed brain sections with antibodies specific to the reporter and cell type specific marker proteins. Specifically, we used anti-GFP antibodies to detect GCaMP5 expression, and quantified co-localization with the astrocyte-specific nuclear marker Sox9 (Sun et al., 2017) to estimate astrocyte specificity. These experiments revealed widespread and strong Cre recombination in about 80% of all Sox9-positive hippocampal and cortical astrocytes (Fig 3A-C). Astrocytes communicate within the astroglial network and with other cell types mainly through intracellular calcium elevations in health and disease (Petzold and Murthy, 2011; Rakers and Petzold, 2017; Reichenbach et al., 2018). Hence, to estimate the usefulness of this Cre line for functional astroglial imaging, we imaged GCaMP5 changes through cortical windows in the cortex and hippocampus under awake and resting conditions as well as under anesthesia (Fig 3D) using two-photon microscopy three weeks after Cre induction. These experiments showed that calcium signals in the cortex in awake mice under resting conditions had a higher amplitude with a longer duration (FDHM) than in the hippocampus of awake and resting mice or in the hippocampus of anesthetized mice (Fig 3E-F).

**Figure 3.**
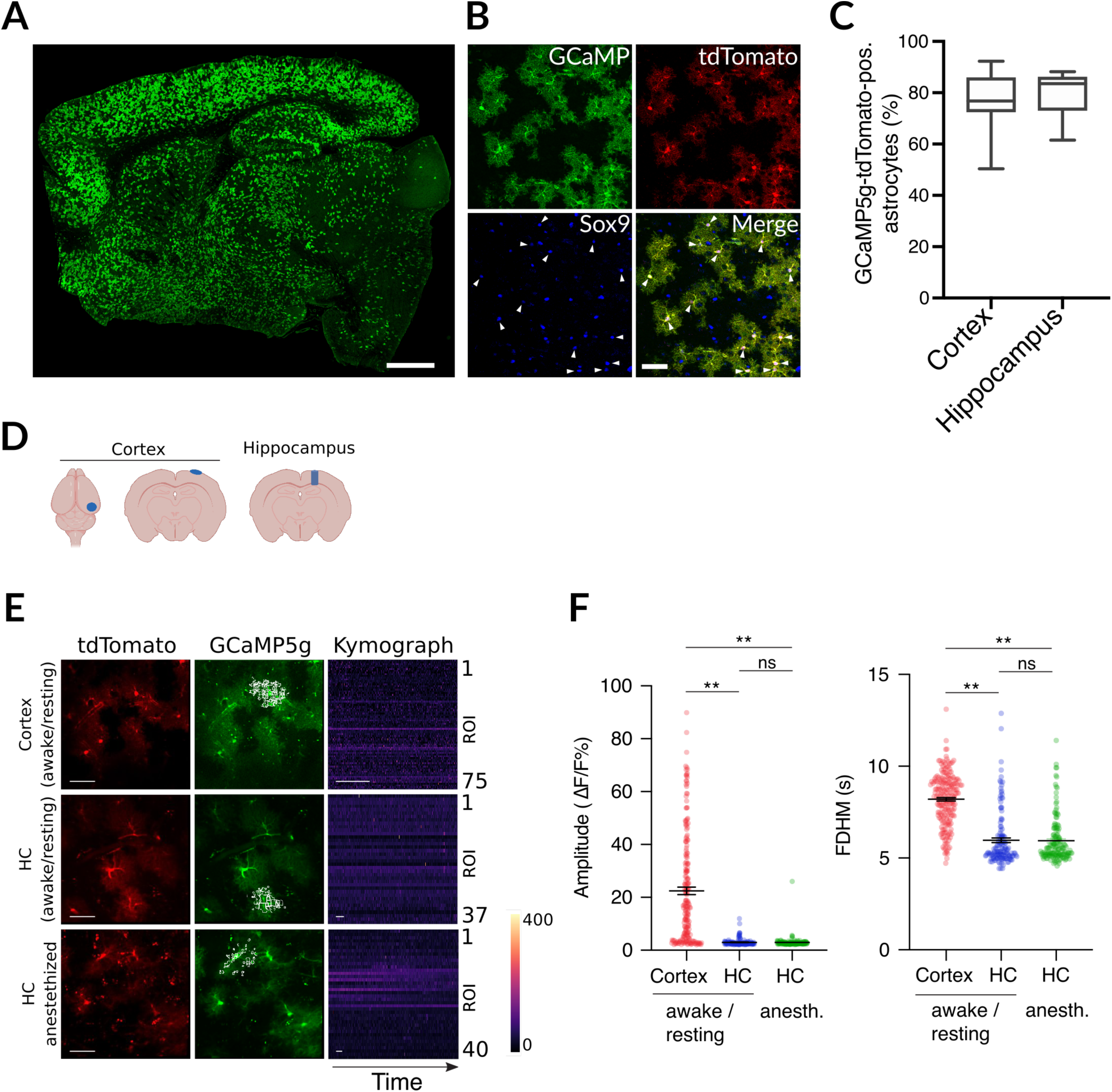
Calcium imaging. (**A**-**B**) Immunofluorescence of Slc1a3-CreERT2::GCaMP5g-tdTomato-loxP using antibodies against GFP and the astroglial marker Sox9 showed widespread and strong Cre recombination specifically in astrocytes (arrowheads; scale bars, 1 mm and 50 µm, respectively; Sox9 nuclear marker was used to identify astrocytes. (**C**) Quantification of recombination efficacy in hippocampus and cortex; reporter expression was normalized to all Sox9-positive cells. (**D**) Cranial windows were implanted over cortex and hippocampus. (**E**) Representative examples of calcium activity under awake/resting conditions in the cortex, and under awake/resting conditions or anesthesia in the hippocampus (HC). No changes were seen in tdTomato fluorescence (scale bars, 50 µm), indicating stable imaging conditions over time, while activity-evoked calcium changes could be detected using GCaMP5g fluorescence. The kymographs represent color-coded calcium changes over time (scale bars, 30 s) in distinct regions of interest (ROI) from individual astrocytes (marked by white boxes in the GCaMP5g images). (**F**) Quantification of calcium activities in the cortex and HC under different conditions revealed that calcium signals in the cortex of awake mice had a higher amplitude and were longer (FDHM, full duration at half-maximum) than in the HC of awake or anesthetized mice (cortex, n = 212 cells from n = 6 mice; HC, n = 141 cells under awake/resting conditions and n = 150 cells under anesthesia from n = 5 mice; Kruskal-Wallis test followed by Dunn’s multiple comparisons test).

### Bergmann glia express markers similar to cerebral astrocytes, but their translatomes are profoundly different

We next sought to characterize the diversity of mRNAs undergoing translation (translatome) in Bergmann glia of the cerebellum and astrocytes of the rest of the brain (hereafter BG and CA, respectively). To this end we crossed Slc1a3-2A-CreERT2 and RiboTag mice (Sanz et al., 2009) congenic for S4 (Rakers et al., 2019). RiboTag mice carry a floxed allele of the endogenous Rpl22 (large subunit ribosomal protein 22) gene that expresses a HA (hemagglutinin) antibody epitope-tagged version of Rpl22 following Cre recombination. CreERT2 was induced with daily intraperitoneal tamoxifen injections for a total of three consecutive days and mice were sacrificed 4 days after the final injection. To evaluate the specificity of recombination, we co-labeled formaldehyde fixed brain sections with an HA antibody for detection of RiboTag and antibodies against marker proteins specific for astrocytes or non-astrocyte cell types. HA positive cells looked like astrocytes and expressed astrocyte marker proteins such as S100ß, Sox9 or GFAP (glial fibrillary acidic protein), but not Osp (oligodendrocyte-specific protein), NeuN (neuronal nuclei), or Iba1 (ionized calcium-binding adaptor molecule 1, microglial marker) (Fig 4). Thus, as observed for GCaMP5, RiboTag expression is activated in the majority of astrocytes, and only astrocytes, by Slc1a3-2A-CreERT2.

**Figure 4.**
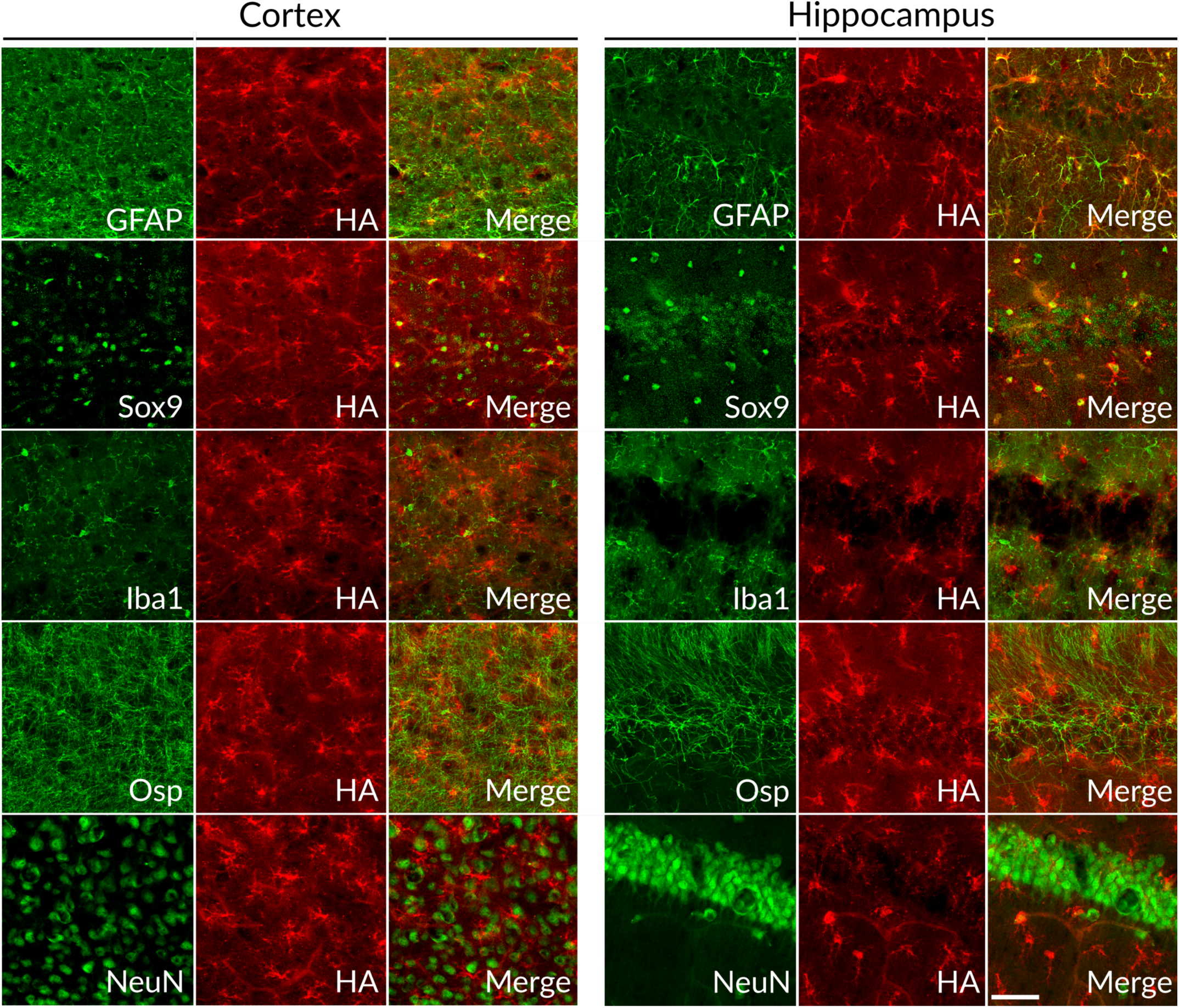
Immunofluorescence staining of Slc1a3-CreERT2::RiboTag mice. Representative images from hippocampus and cortex of 7 weeks old mice. For each marker, HA (hemagglutinin) co-staining (RiboTag, red) was used to mark recombined cells; GFAP – glial fibrillary acidic protein, Sox9 - SRY-Box Transcription Factor 9 (astroglial marker), Iba1 - ionized calcium binding adaptor molecule (microglial marker), Osp – oligodendrocyte surface protein, NeuN – neuronal nuclei. Scalebar = 50μm.

We next studied the translatome of astrocytes from BG and CA by capturing HA-tagged ribosomes with immunoprecipitation (IP) and analyzing the captured mRNAs with next generation sequencing (NGS). We first performed a principal component analysis which revealed that replicates of a given sample type clustered tightly together, indicating the IPs were reproducible, but the different types of samples were quite different (Fig S3). The first principal component separated both RiboTag IPs from their corresponding total RNA samples in the same direction and to a similar extent (Fig S3), indicating RiboTag had captured a population of mRNAs very different from the input RNA. The second principal component separated the samples based on the brain region to the same extent for input and IP, indicating that the translatomes of BG and CA are very different. We then tested if the differentially expressed genes were sufficient to reveal the identity of the cells from which the mRNAs were captured. Gene set enrichment analysis (GSEA) demonstrated that the top 10 pathways enriched in IPs from both regions involved processes of metabolism, catabolism, and detoxification, known core functions of astrocytes (Fig 5A, B). In contrast, the top 10 pathways depleted from IPs from both regions involved processes related to neuronal activities and synaptic structures (Fig 5A, B). These results indicate that an unbiased analysis was sufficient to establish that the IP-captured samples were enriched for mRNAs expressed by astrocytes and depleted for mRNAs expressed by neurons.

**Figure 5.**
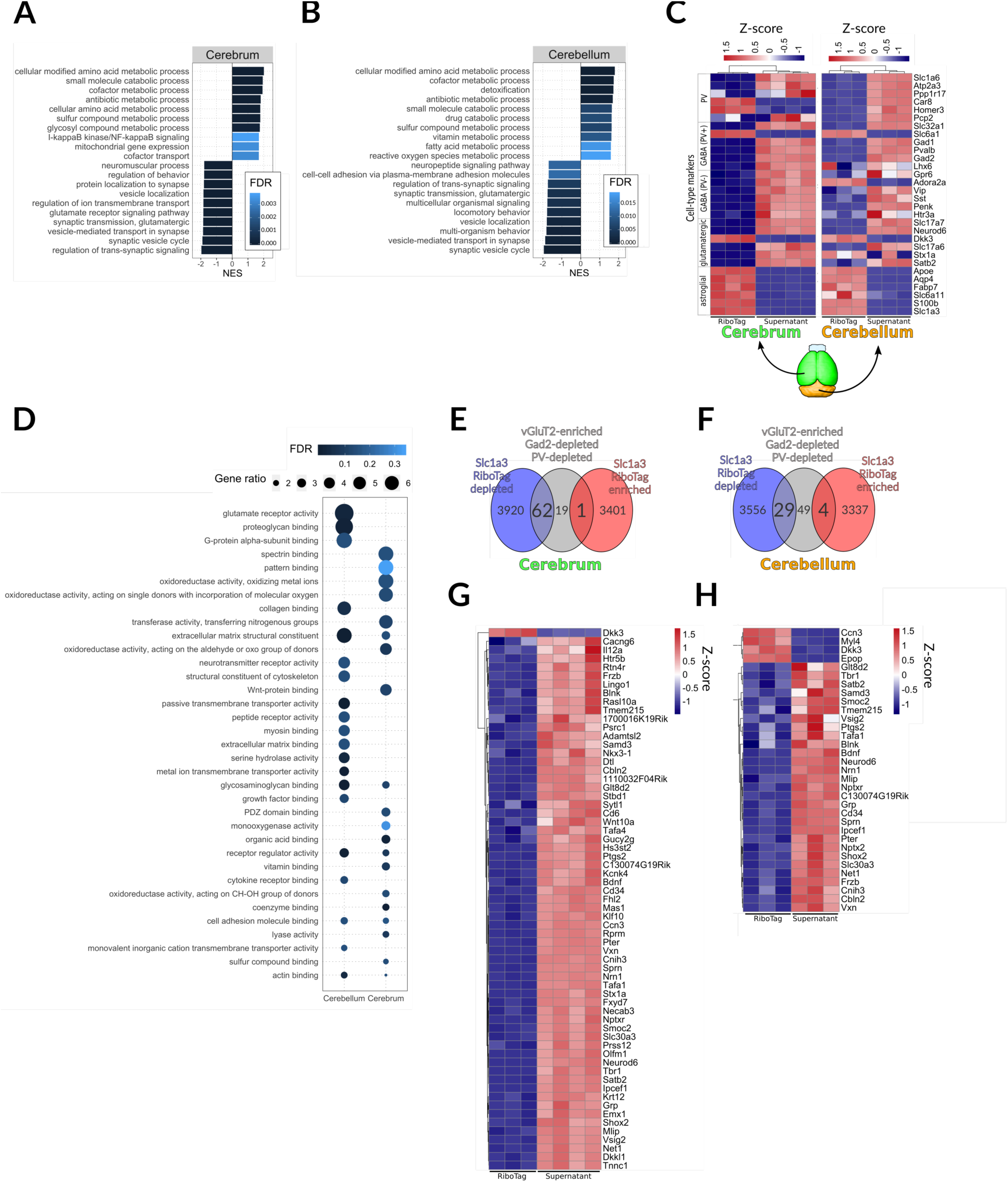
Assessment of RiboTag analysis specificity. (**B, C**) Gene set enrichment analysis (GSEA) of RiboTag IP in cerebrum (**B**) and cerebellum (**C**); negatively and positively enriched genes correspond to gene sets functionally related to neuronal and glial function, respectively; analysis was done using WebGestalt (Liao et al., 2019) using molecular function GO terms, genes were ranked as described in the methods section and top 10 hits for each region were shown. (**D**) Heatmap showing relative distribution of RiboTag RPKM values for genes for cell type-enriched mRNAs that we previously used to validate cell type specificity of gene expression study in Tagger mice (Kaczmarczyk et al., 2019). Each column represents one biological replicate. Z-score was calculated as follows: Z = (x − meanrow(x))/SD(row), where SD is standard deviation. (**E, F**) Venn diagrams showing overlaps of significantly enriched (red circle, LFC > 1, FDR < 0.05) and depleted (blue circle, LFC < −1, FDR < 0.05) genes in Slc1a3-2A-CreERT2::RiboTag with genes enriched in glutamatergic neurons in vGluT2-Cre::Tagger mice. Grey circle contains genes that are enriched in vGluT2+ cells (LFC > 1, FDR < 0.05), and simultaneously not enriched in PV+ neurons and Gad+ neurons (LFC < 0, FDR < 0.05 for both cell types). Note that in case of our Tagger analysis whole brain was used, hence the expected overlap (circle intersections; genes enriched in vGluT2 should be depleted in astroglia and vice versa) is more pronounced for the cerebrum (more similar to Tagger samples) (**E**) than for the cerebellum (**F**). (**G, H**) Heatmaps showing genes from the intersections of the Venn diagrams in E and F.

We further validated the cell type specificity of the captured mRNAs by analyzing a list of marker genes generated with RiboTag data for a previous study that comprises many known cell type markers (Kaczmarczyk et al., 2019). As expected, markers specific for astrocytes (Apoe, Aqp4, Fabp7, Slc6a11, S100b and Slc1a3) were highly enriched in RiboTag IP samples relative to total RNA, whereas markers for a wide diversity of neurons were depleted (Fig 5C), with some notable exceptions. Adora2a (adenosine receptor 2a) is commonly considered a marker for astrocytes but is also a strong marker for GABAergic neurons in the striatum. Consistent with this, Adora2a was enriched in RiboTag samples from BG but depleted from RiboTag fractions from CA, where the striatum is located (Fig 5C). Slc1a6 is a GABA transporter and was included in our list as a marker of GABAergic neurons. However, in the current study it appeared to be preferentially expressed by BG and CA, suggesting a more prominent role for GABA uptake by Slc1a6 for astrocytes than neurons. Likewise, Gria1 and Gria4, both involved in AMPA signaling in Bergmann glia (Ben Haim and Rowitch, 2017; Matsui et al., 2005), were enriched in RiboTag samples from BG but not CA. We then compared these data to a larger manually curated list based on literature with the same overall result that astroglial markers were enriched and markers of other cell types were depleted (Fig S4). We then did gene ontology analysis on the list of genes significantly different (LFC > 1, FDR < 0.05) between BG and CA. Consistent with previous observations of expression of glutamate receptors by BG, the most enriched molecular function gene ontology category in BG was “glutamate receptor signaling” (Fig 5D). That result motivated us to ask how similar the translatomes of BG are to those of glutamatergic neurons. Employing previous data (Kaczmarczyk et al., 2019), we curated a list of mRNAs that are specifically enriched in glutamatergic neurons compared to GABAergic and parvalbumin neurons and determined if this selection overlapped with mRNAs that were either enriched or depleted in astrocyte translatomes. We found 82 genes were specific for glutamatergic neurons, of which only 1 was enriched in CA whereas 4 were enriched in BG (Fig 5E-H). In contrast, 62 and 29 genes were depleted from CA and BG translatomes, respectively (Fig 5E-H), indicating that expression of glutamatergic-related signaling systems is very limited in CA but present in BG, and suggesting that glutamatergic signaling is a functional difference between the two astrocyte types (Saab et al., 2012). Overall, these analyses supported the immunofluorescence study indicating the Slc1a3-2A-CreERT2 line specifically activated RiboTag in astrocytes. Furthermore, they highlighted that the translatomes of BG and CA are remarkably different.

We then explored these data for new biological insights. To this end we focused on genes known to be involved in neurodegenerative diseases (NDs). NDs result in the slow degeneration of the central nervous system. The late onset nature of NDs is especially remarkable for cases of inherited neurodegenerative diseases where the disease-causing protein is expressed throughout life. They are marked by death of neurons and morphological changes of astrocytes, sometimes becoming dysfunctional (Phatnani and Maniatis, 2015). We therefore examined the expression pattern of a manually curated list of the most prominent genes linked to neurodegenerative diseases, and observed many to be unequally expressed across brain regions and cell types (Fig 6). The mRNA from the causative gene in prion diseases, Prnp, displayed a relative enrichment in astrocytes in the cerebrum (Fig 6) consistent with a previous report (Jackson et al., 2014). Interestingly, three mRNAs linked to polyglutamine diseases, Sca1, Sca3 and Htt, appeared to be depleted from astrocytes and therefore primarily neuronally expressed, consistent with neurons preferentially producing the disease related protein aggregates. Also, HPRT, a housekeeping enzyme, was previously used as an ectopic carrier of polyglutamine and similarly demonstrated a preferential expression in non-astroglial cells, potentially explaining the aggregation of polyglutamine-HPRT specifically in neurons and the severe phenotype observed in that model (Jackson et al., 2003; Ordway et al., 1997). In contrast to these examples of monogenic diseases, amyo trophic lateral sclerosis (ALS), Parkinson’s (PD) and Alzheimer’s (AD) diseases are linked to multiple genes. Interestingly, for a given disease, some genes appear to be preferentially expressed by CA and/or BG whereas other genes appeared to be preferentially expressed by other cells. One striking example includes amyloid precursor protein (App) and presenilin 1 (Psen1), both linked to AD. Psen1 is the key component of gamma secretase, a membrane localized protein cleaving complex involved in notch pathway signaling and the generation of toxic fragments of App. Surprisingly, App was depleted in IPs and presumably preferentially expressed by neurons, whereas Psen1 was highly enriched specifically in BG, indicating that the expression pattern of App is determined by factors besides Psen1. PD related genes also displayed contradictory expression patterns where alpha synuclein (Snca, known to be neuron specific) was depleted in CA, while Prkn, Park7 and Pink1 enriched in astrocytes of both regions. Likewise, of genes associated with ALS, Fus was enriched in CA whereas C9orf72 was depleted in BG. Thus, multiple genes causative of the same ND often have remarkably different expression patterns. Finally, we noted that the mRNA encoding RiboTag, Rpl22, appeared to be relatively enriched in IP fractions. This prompted us to examine the expression pattern of other ribosomal proteins.

**Figure 6.**
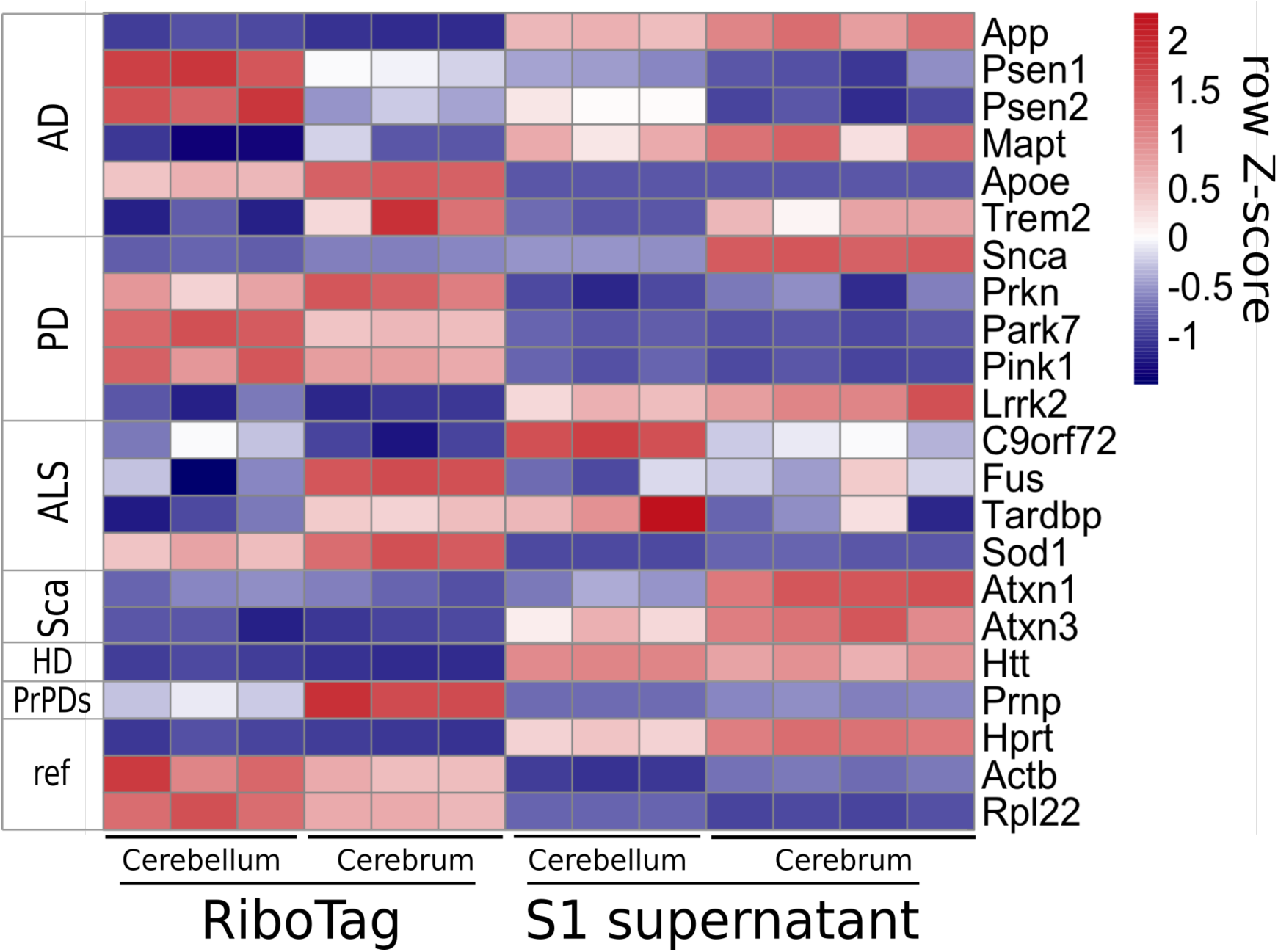
Expression of ND-related genes across the cerebellum and the rest of the brain in astroglia (RiboTag) and total tissue (input samples).

Ribosomes are large protein synthesizing machines highly conserved across all kingdoms of life, consisting of four ribosomal RNAs and approximately 80 proteins in eukaryotes. In text books they are portrayed as being uniform and primarily differing only in respect to regulatory phosphorylation. However, the hypothesis that ribosome heterogeneity may function as a new layer of gene expression control is steadily gaining attention (Emmott et al., 2019b; Genuth and Barna, 2018). We therefore examined the expression profile of all the core ribosome proteins. Surprisingly, we observed a non-uniform distribution of mRNAs encoding ribosomal proteins (Fig 7A). The vast majority of ribosome protein encoding mRNAs were more highly represented in IP fractions, suggesting that the synthesis of ribosomes commands a higher proportion of translational output for astrocytes compared to other brain cells. Interestingly, some appeared to be enriched specifically in BG (e.g. Rps8 and Rpl39), whereas others (e.g. Rps6, Rpl30 and Rpl10a) appeared to be depleted in BG (Fig 7A). Since ribosome protein levels are not always correlated with their mRNA levels (Emmott et al., 2019a), we stained brain sections with antibodies against various ribosome proteins. We used Slc1a3-RiboTag brain sections so the RiboTag protein could function as a convenient histological marker for BG, the cells our RiboTag samples were specifically captured from, and the rat derived HA antibody we commonly use could complement the available ribosomal protein-specific antibodies that are typically from rabbit. These experiments revealed that Rps21 was prominently expressed throughout the cerebellum, especially in BG but less prominent in Purkinje neurons (Fig 7B). In contrast, Rpl26 was prominently expressed by Purkinje neurons but less so by other cells including BG (Fig 7B). This striking difference in distribution of the proteins was unexpected as the mRNAs suggested they share a similar expression pattern (Fig 7A) and indicates that factors beyond mRNA levels control the distribution of ribosome proteins. In summary, these translatome data provide a resource to study the characteristics of BG that make them unique.

**Figure 7.**
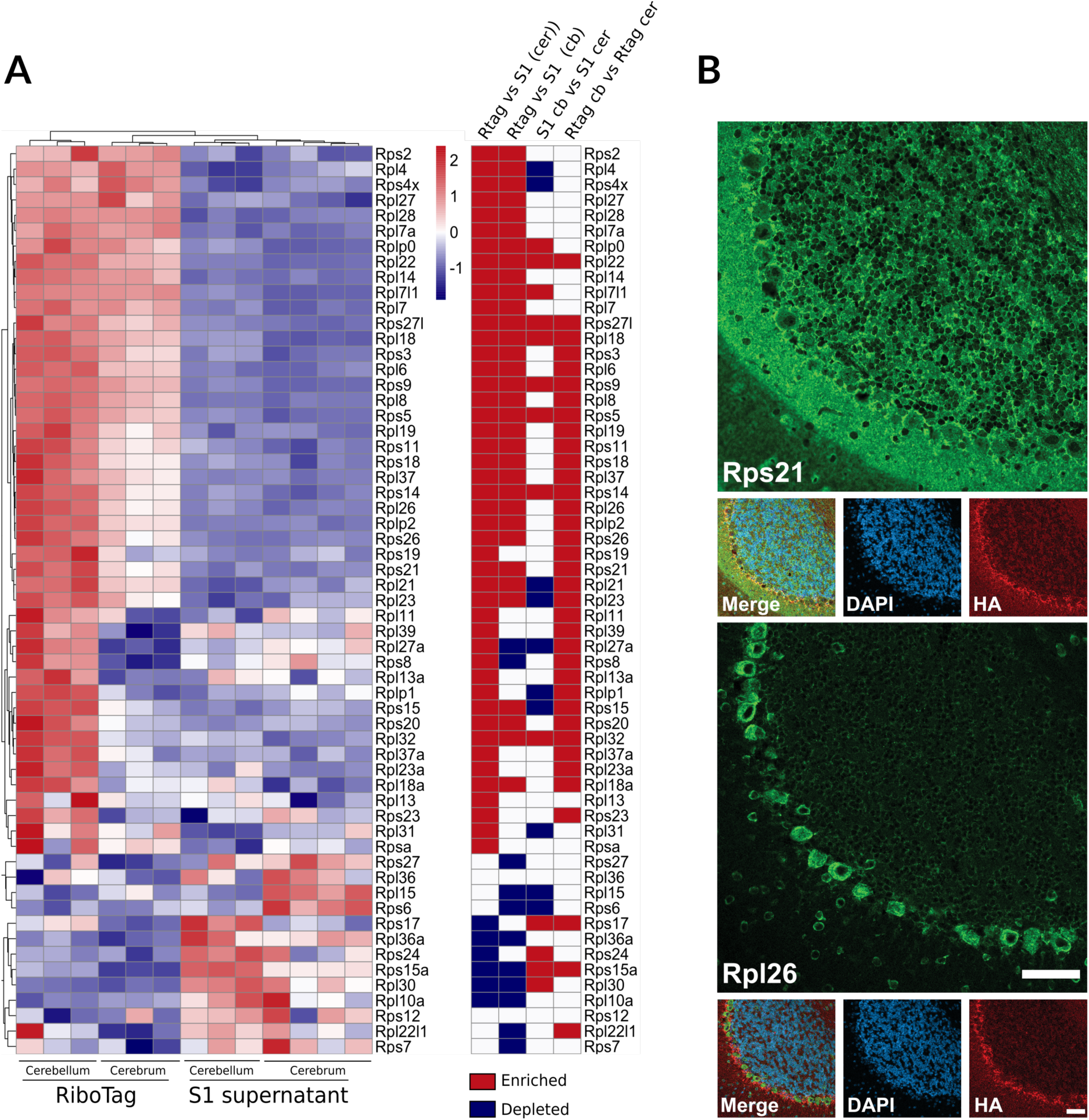
(**A**) Heatmaps of FPKM values for significantly (FDR < 0.05) enriched and depleted (RiboTag vs input) ribosomal proteins in cerebellum and cerebrum. (**B**) Immunofluorescence show disparity between ribosomal subunits Rps21 and Rpl26 in the cerebellum.

## Discussion

Minimization of genetic variability by inbreeding has become a gold standard in research using mouse models. Selection of mice from highly uniform populations increases the statistical power, thereby making experiments more sensitive and efficient, enabling a reduction in the number of replicates and mouse lives needed. The desire for using inbred mouse lines has led to many being available exclusively on a B6 background. However, the high inter-individual variability we observed in congenic B6 mice (Jackson et al., 2009; Jackson et al., 2013) and here in inbred B6 mice could confound experimental results, especially those involving high content gene expression analyses. For this reason, it is worthwhile considering alternative genetic backgrounds. Our results should not, however, be interpreted to suggest that B6 mice are abnormal. In fact, high activity and tendency to explore, if interpreted as eagerness to escape captivity, is how wild mice ought to behave. In this sense, S4 mice might be considered abnormal due to their docile character and low activity levels. Although they were historically bred to study testicular cancer (Stevens and Little, 1954), we are aware of only a single case of spontaneous testicular tumor formation during several years of work with multiple lines in this genetic background, often aged up to two years. Importantly, most embryonic stem cell lines used for the generation of gene-targeted mice are derived from 129 strains, especially S4, and the S4 strain is a robust breeder, so it is a logical complement to B6 strains. Therefore, the highly homogeneous nature of S4 is desirable for studies of gene expression which prompted us to establish several mouse lines in this alternative background (Fig S2). A mouse model of AD is already available in the S4 background (https://www.jax.org/strain/031988) and we have also backcrossed to S4 a knock-in mouse model of HD (Lin et al., 2001).

### Precise targeting of astroglia is challenging

Although not directly involved in electrical signaling, astrocytes are essential for normal brain function and outnumber neurons in higher mammals. They play crucial roles in synaptic plasticity (Singh and Abraham, 2017), removing extracellular waste (Xie et al., 2013), and actively participate in neural processes controlling behavior (Oliveira et al., 2015). Disruption of astrocyte function contributes to a variety of diseases (Li et al., 2019; Liddelow and Sofroniew, 2019; Sofroniew and Vinters, 2010; Tian et al., 2005). Therefore, a steady need for tools to access astrocytes *in vivo* exists. However, several features of astrocytes make them notoriously difficult to genetically target (Garcia et al., 2004; Hirrlinger et al., 2006; Khakh and Sofroniew, 2015). First, they make less total protein, and the half-lives of their proteins are typically shorter than in neurons (Dorrbaum et al., 2018). As a result, the native levels of effector proteins expressed in astrocytes are often insufficient for completing the desired function. Many of the earliest lines developed were based on constitutive activity of Cre from mouse or human GFAP promoter elements, which have been reported to target neurons as well (Xie et al., 2015; Zhuo et al., 2001). The flaws of this approach have since been thoroughly scrutinized (Sloan and Barres, 2014). Many marker genes specific for adult astrocytes are also expressed in NPCs during development (Ferenczy et al., 2013; Foo and Dougherty, 2013), many of which become neurons. This necessitates the employment of inducible strategies, such as CreERT2, although they can be inefficient or hard to tune (Chow et al., 2008; Kristianto et al., 2017). The Slc1a3 gene was used previously to drive Cre using a BAC method whereby a large region of genomic DNA encompassing an entire gene is modified to carry coding sequence for Cre and then returned to the genome where it lands in a random location. BAC transgenic mice have important strengths, such as being easier to develop and can provide much higher expression levels than the native gene. However, the recombination efficiency and specificity of a BAC-GLAST-Cre line (using Slc1a3’s alternative name) was variable depending on the founder line employed (Slezak et al., 2007). Since each line likely carries the BAC in a different genomic location and at various copy numbers, variable activity was likely due in part to the use of a BAC approach where native elements of the genomic integration site can compete with the regulatory elements carried by the transgene. Moreover, some BAC-GLAST-CreERT2 mice showed significant recombination in neurons, in spite of utilizing the same promoter elements, possibly due to multiple integrations, or integrations in the vicinity of neuronal enhancers and, consequently ectopic expression of Cre (Kristianto et al., 2017). Notably, that line also targeted NPCs. We did not check if this is the case for our Slc1a3-2A-CreERT2 knock-in line, but since NPCs express Slc1a3 we consider this plausible. Ectopic labeling of neurons due to activation of NPCs might be avoided by limiting the time window between activation and sacrifice. We also did not test the specificity of our line in the peripheral nervous system or other organs. Efforts to do so in the future will be well placed.

Another potential problem with BAC transgenics is that the invading transgene does more than express CreERT2. For example, besides inserting large (often hundreds of kilobases) fragments of ectopic DNA that can interrupt native genomic neighborhoods (Kaczmarczyk and Jackson, 2015), it is not uncommon for large deletions to also occur at the site of integration (Goodwin et al., 2019). Moreover, BACs typically carry additional genetic elements. For example, in BAC Aldh1l1-CreERT2 mice the transgene appears to also carry genes encoding a protein (Slc41a3), a miRNA, and a long non-coding RNA (Srinivasan et al., 2016). Slc41a3 is a magnesium transporter highly expressed in cerebellar Purkinje cells. Since magnesium is a modulator of neurotransmitter receptors (Gentry et al., 2015; Hu et al., 2020) and cofactors and/or allosteric modulators of enzymes (Bachelard, 1971; Kirkland et al., 2018), overexpression of Slc41a3 might alter brain homeostasis. The effects of overexpressing the miRNA and long non-coding RNA are more difficult to predict but should not be ignored. Fortunately, problems associated with BACs can be avoided by using gene targeting methods. A gene-targeted mouse line using the Slc1a3 gene was developed before ours (Mori et al., 2006). That line resulted in the complete inactivation of the host gene, whereas ours was designed to leave expression unscathed. Although Slc1a3 knock-out mice are viable, they display some differences from wild types (Karlsson et al., 2009). We attempted to avoid this through use of the 2A system, whereby expression of Slc1a3 and CreERT2 would be expressed from a single open reading frame. However, the residual 2A peptide on Slc1a3 appeared to reduce the steady state level. Furthermore, we have not determined if Slc1a3 protein carrying the 2A peptide is functional. This unexpected observation may call for caution when applying the 2A peptide method to express two proteins from a single mRNA. However, any phenotypic effects it might have for Slc1a3-2A-CreERT2 mice will likely be negligible. We maintain the line as homozygotes, we have observed no abnormalities, and they behave, live and reproduce like wild-type 129S4 counterparts.

In our pilot studies using RNA ISH, compared to Gja1 and Slc1A3, Aldh1l1 was lowly expressed, consistent with data presented on the Allen Brain Atlas (https://mouse.brain-map.org). Nonetheless, the Aldh1l1 gene was successfully used in the generation of both a BAC CreERT2 line (described above) and a knock-in CreERT2 mouse line using a 2A approach like ours (Hu et al., 2019). Aldh1L1-T2A-CreERT2 mice were shown to have perfect specificity and very high efficiency, 50 to 80%, depending on the region and induction protocol (Hu et al., 2019). The gene-targeted Slc1a3 Cre-ERT2 line mentioned above (Mori et al., 2006) was tested in parallel and both lines had very favorable and similar performance (Hu et al., 2019), consistent with the data we present here. It has been reported that a given Cre line can yield very different recombination efficiencies of floxed alleles at different loci, likely due to differences in expression intensity and therefore chromatin states (Dixon et al., 2004). Our work employed reporter lines at two different genomic loci. PC::G5-tdT was inserted 1 kb downstream of the Polr2a gene whereas RiboTag was built into the Rpl22 gene. Both are naturally highly and ubiquitously expressed genes which may have benefited our study with good induction for both. Indeed, the evaluation of the Aldh1l1-T2A-CreERT2 knock-in line employed a 7-day induction protocol followed by 2 weeks of expression whereas we employed milder induction strategies that yielded similar induction rates. Thus, Aldh1l1-T2A-CreERT2 knock-in mice may yield an even higher efficiency if paired with an alternative reporter line. Unfortunately, the genetic background of the Aldh1L1-T2A-CreERT2 knock-in line was not clearly described.

### Ribotag reveals curious cases of contradictory mRNA distribution in the brain

To validate the S4 Slc1a3-2A-CreERT2 line we developed RiboTag translatome data from two regions of the mouse brain. Besides establishing that RiboTag was activated specifically in astroglia, these data served two purposes. First, they offered a comparison of translatomes from Slc1a3-expressing glia from two brain regions, which in the case of cerebellum, are specifically BG. The comparison of translatomes of BG to CA revealed that some genes thought to be markers for astrocytes or non-astrocytes are selective for one brain region but not the other, expanding the concept that astrocytes are highly diverse. Second, the RiboTag data resulted in two surprising findings. First, we found that genes causative of a single ND had disparate expression patterns. For example, Psen1 is important for processing App into disease forms that cause AD, but their expression patterns are different (Fig 6). Likewise, genes involved in PD and ALS also demonstrated contradictory expression patterns (Fig 6). Second, our data may be interpreted as supporting the hypothesis of ribosomal specialization (Xue and Barna, 2012). Although it is controversial (Ferretti and Karbstein, 2019), the concept of specialized ribosomes posits that the stoichiometry of ribosome proteins on ribosomes results in preferential translation of a subset of mRNAs in the translatome (Shi et al., 2017; Yamada et al., 2019). While our observations of non-stoichiometric expression do not provide conclusive proof for ribosomal specialization, they do suggest that conditions needed for specialized ribosomes exist in the brain.

S4 Slc1a3-2A-CreERT2 mice are available at the European Mouse Mutant Archive <u>EM:11807</u> and the RiboTag gene expression data described here can be accessed at https://jaws.research.liu.se/resources.html

## Materials and Methods

### Ethical statement

Ethical permissions for this work were granted by the Landesamt für Natur, Umwelt und Verbraucherschutz Nordrhein-Westfalen, permission #84–02.04.2012.A192, #84– 02.04.2017.A016 and #84-02.04.2016.A442, or the Massachusetts Institute of Technology Committee on Animal Care #0705-044-08. All experimental procedures were performed in accordance with the internal regulations of the DZNE or MIT.

### Mouse line generation

Targeting vector was constructed in the pBlueScript backbone using combination molecular biology methods and gene synthesis. Homology regions were PCR-amplified from 129S4 genomic DNA. Cre-ERT2 sequence was pCAG-CreERT2 was a gift from Connie Cepko (RRID: Addgene_14797). The targeting vector is available at Addgene (RRID: Addgene_129409). Gene targeting was done in J1 ES cells (ATCC) as described previously (Kaczmarczyk et al., 2016). The stop codon of Slc1a3 was targeted with Cas9 by the double nickase strategy (Ran et al., 2013), using the pX335 construct (a gift from Feng Zhang, RRID: Addgene_42335), bearing <u>G</u>TT CTC TGT CCG TCT ACA TCT and <u>G</u>CT TTC TTA AGC ACC AA GTG T sgRNA protospacers. Targeting was verified by PCR using LongAmp HotStart polymerase (NEB) using primers listed in Table S1.

Gene targeted cells were injected into C57Bl/6N blastocysts using established methods. A male chimera transmitting the ES cell genome to progeny was identified by first breeding to C57Bl/6N females, and subsequently bred to females expressing Flpo recombinase (Jax line #007844) (Raymond and Soriano, 2007) to simultaneously remove the selection cassette and establish the line in 129S4 background. This F1 generation was bred two more times to 129S4 mice to remove the Flpo transgene and any spontaneous mutations arising from culturing or storage of the ES cells. RiboTag mice were acquired from Jackson labs (Jax line #011029), with a predominately B6 background. They were backcrossed to S4 for 10 generations, periodically guided by SNP analysis for choosing breeders. A final SNP genotyping analysis (Envigo RMS, Inc, Indianapolis, USA) measuring 347 SNPs that discriminate S4 from B6 revealed only 1 residual B6 SNP, indicating the genetic background of the line is approximately 99.7 % S4. For backcrossing work, 129S4 mice were obtained from Jackson labs (Jax line# 009104). For behavioral experiments, wild type B6 (C57Bl/6NTac) mice were acquired from Taconic and wild type 129S4 mice were acquired from the Whitehead Institute/Massachusetts Institute of Technology, where this strain originated, and both strains were bred in house less than 5 generations prior to inclusion in experiments.

### Automated mouse behavioral analysis (AMBA)

Video recordings were performed as described previously at 6, 8, 10, 12, 14, 16, 18, and 20 months of age on female mice (Jackson et al., 2009; Jackson et al., 2013). Since video recordings must be of mice in individual cages and returning males into group housing may induce fighting, males were excluded from the study. Videos were analyzed with Home Cage Scan software (CleverSys Inc.) followed by a customized downstream pipeline (Jackson et al., 2009; Steele et al., 2007). For most mice weight was measured monthly. In a first cohort of mice weight measurements were initially stopped when it was noticed that some B6 mice were highly variable. In the second cohort mice were weighed at 18 months. Violin plots were generated using ggplot2 (https://ggplot2.tidyverse.org). Horizontal lines represent medians.

### Tamoxifen injections

To induce expression of the GCaMP5g-tdTomato reporter, mice were injected i.p. with 5 μl/g body mass with an emulsion of 20 mg/ml tamoxifen (Sigma, #T5648) in sunflower oil:ethanol mix (10% ethanol). Sunflower oil was from Sigma (#47123). Three months old mice were injected for five consecutive days, followed by imaging 3 weeks later. To induce expression of the RiboTag reporter, mice were injected with 10 μl/g body mass with an emulsion of 10 mg/ml tamoxifen (Sigma, #T5648) in sunflower oil:ethanol mix (10% ethanol). Injections were done daily for three total injections, and mice were sacrificed 4 days after the final injection at approximately 2 months of age for RiboTag studies.

### Cranial window preparation

Hippocampal window surgery was performed 4 weeks before imaging as described previously (Reichenbach et al., 2018). Mice were anesthetized with isoflurane (induction, 3%; maintenance, 1–1.5% vol/vol; Virbac), and body temperature was maintained with a heating pad (37°C). Mice received buprenorphine (0.1 mg/kg; Reckitt Benckiser), dexamethasone (0.2 mg/kg; Sigma Aldrich #D1159) and cefotaxime (2 g/kg; Fisher Scientific #15219946). After fixation in a stereotaxic frame, the skin was removed under sterile conditions and a craniotomy (diameter, 3 mm) above the right somatosensory cortex (coordinates: AP –1.9 and ML +1.25 relative to bregma) was created with a dental drill. The dura was removed, and the somatosensory cortex was removed with a 21G needle attached to a 20-ml syringe with a flexible tube. When the external capsule of the hippocampus was reached, the alveus was carefully exposed using a 27G needle. Subsequently, a metal tube (diameter, 3 mm; height, 1.5 mm) sealed with a glass coverslip (diameter, 3 mm) was inserted, and the upper tube edge was glued to the skull bone using dental cement. The remaining exposed surface of the skull bone was sealed with dental cement, and a custom-made metal bar was glued next to the metal tube. Mice received buprenorphine (0.1 mg/kg) for 3 d after surgery.

For cortical window preparation, mice were anesthetized with isofluran (induction, 3%; maintenance, 1–1.5% vol/vol), and body temperature was maintained with a heating pad (37°C). Mice received buprenorphine (0.1 mg/kg), dexamethasone (0.2 mg/kg) and cefotaxime (2 g/kg). After the mice were fixed in a stereotactic frame, the scalp was removed and a craniotomy (diameter, 3 mm) was created above the right somatosensory cortex using a dental drill. The surface was rinsed with sterile saline, a coverslip (diameter, 3 mm) was inserted and glued with dental cement, and a headpost (Luigs & Neumann) was glued adjacent to the cortical window. Mice received buprenorphine (0.1 mg/kg) for 3 d after surgery. Schematic images of cranial windows were prepared using BioRender (https://biorender.com).

### *In vivo* two-photon microscopy

Mice implanted with hippocampal windows were imaged using an upright two-photon microscope (Trim ScopeII; La Vision) with a 16x objective (NA 0.8, Nikon LWD16x) and three non-descanned detectors with two bandpass filters (617/73, 525/50 nm) and one long-pass filter (550 nm). Mice were imaged anesthetized (with isoflurane; 1-1.5% vol/vol) and awake on a treadmill (Luigs & Neumann). For hippocampal imaging, the same group of mice was imaged under awake and anesthetized conditions. Fluorophores were excited at 920 nm using a Titan Sapphire (Ti:Sa) laser (Chameleon Ultra II; Coherent; 140-fs pulse width, 80-MHz repetition). XY time-lapse series of astroglial calcium activity (256 × 256 μm; 256 px; pixel dwell time, 1.62 μs) were subsequently recorded for 10 min at 3.61 Hz at a depth of 100– 200 μm beneath the hippocampal surface. Mice implanted with cortical windows were imaged awake on a treadmill using an upright two-photon microscope (SP8 DIVE (Deep *In Vivo* Imager); Leica) with a 16x objective (NA 0.8, Nikon LWD 16x) and hybrid detectors (HyD) with two filters (500/550, 560/620 nm). Fluorophores were excited at 920 nm using a Spectra Physics InSight DS+ laser. XY time-lapse series of the astroglial calcium activity (256.28 × 256.28 μm; 256 px; pixel dwell time, 975 ns) were recorded for 10 min at 3.82 Hz at a depth of 100-200 μm beneath the cortex surface. Laser power below the objective was kept between 20 and 40 mW to minimize laser-induced artifacts and phototoxicity.

### Calcium imaging data analysis

Calcium imaging data were stabilized using a custom-written Lucas-Kanade algorithm (Guizar-Sicairos et al., 2008) in Matlab R2018a (MathWorks). Regions of interests (ROIs) representing astroglial microdomains were defined by fluorescence changes over time in GCaMP5g-expressing astrocytes using a custom-written macro (National Institutes of Health) in ImageJ 1.50i (Agarwal et al., 2017). Time-lapse data for each ROI were normalized, smoothed, and peak candidates were detected with a hard threshold. Detection and classification of fluorescence peaks over time was performed with a custom-written algorithm in Python. Mean fluorescence data were first normalized by a robust z-score calculated per ROI over the whole time-lapse series. Normalized data were then smoothed with a Gaussian filter, and all maxima above the threshold were selected as peak candidates. Peak candidates were defined by its ROI and the timepoint of peak maximum. Peak amplitude and full duration at half maximum (FDHM) were determined for each peak candidate. Each time-lapse series was plotted together with the respective video file for visual inspection and verification.

### RNA-seq and data analysis

RiboTag-purified and total RNA samples were sequences at Atlas Biolabs. Library was prepared with TruSeq Stranded mRNA protocol and samples were indexed using TruSeq RNA CD Index Plate. Libraries were QC-ed and quantified on TapeStation system (Agilent) and 50 bp single reads were obtained from HiSeq2500 sequencer (Illumina). Demultiplexing was done using standard Illumina pipeline and reads were QC-ed using FastQC. Reference (GENCODE GRCm38) was indexed, and the reads were aligned with BBmap using the following parameters: -Xmx30g in=in.fq out=out.bam qtrim=t usequality=t minaveragequality=0 local=f strictmaxindel=f xstag=us maxindel=100000 intronlen=10 ambig=toss threads=8. BAM files were indexed and sorted with Samtools v1.9 and exon counts were obtained with summarizeOverlaps() R function. RPKM values and gene enrichments/depletions were computed using DEseq2 R package (Love et al., 2014). Heatmaps were generated using pheatmap() R package. Statistical significance of gene list overlaps (Venn diagrams) was computed with hypergeometric test using phyper() R function. Gene ontology analysis was done with WebGestalt (Liao et al., 2019). Complete RPKM data is available at https://jaws.research.liu.se/resources.html and at GEO database (GEO GSE145484, https://tinyurl.com/s8gtkur).

### Immunohistochemistry

**Ribosomal proteins (Fig 7)** At 6 weeks weeks of age mice were sacrificed and brains removed and emersion fixed in 10% formalin for two days. Brains were then processed through zylenes and paraffin and placed in cassettes. Cassettes were cut into 4 µm thick sections. Sections were dewaxed in histolab clear (Histolab #14250) and rehydrated in graded dilutions of ethanol, 100-0% (each 5 min). Epitope retrieval was performed with a steamer in 0.01M citrate buffer, pH8, for 20 min and cooled down for 15 min at RT before incubated in dH_2_O for 5 min. Auto fluorescence was quenched using the TrueBlack Lipofuscin Autofluorescence Quencher (Biotium #23007) according to the manufacturer’s protocol. Sections were incubated in blocking buffer (PBS with 2.5% NHS) for 30 min prior to addition of primary antibody solution for 2 h at RT. Double stains were performed using a mix of the two primary antibodies in blocking buffer simultaneously. Then, sections were washed in PBS for 3 × 5 min. After the last wash step secondary antibodies diluted in blocking buffer were added for additional 30 min incubation, RT. A second wash with PBS for 3 × 5 min, RT, was conducted prior to the use of a second auto fluorescence quenching kit (Vector TrueVIEW autofluorescence quenching kit, Vector Laboratories, # SP-8400). Finally, sections were washed in PBS for 5 min, incubated with DAPI (conc. 0.1 µg/ml) for 5 min and washed in PBS for 5 min. Mounting was performed using Vectashield vibrance antifade mounting media (Vector Laboratories, #H-1700). Sections were imaged on the confocal microscope LSM 800 (Zeiss) using a 20x objective. Primary antibodies: HA-tag, 1:100 (Roche, RRID: AB_390919), RPS21 1:100 (Bethyl, RRID: AB_2631466), RPL26 1:100 (Cell Signaling, RRID: AB_10698750). Secondary antibodies (each used at 1:500): Alexa Fluor 647-goat-anti-rat (Thermo Fisher), Alexa Fluor 488-donkey-anti-rabbit (Jackson ImmunoResearch, RRID: AB_2313584). **GCaMP5g, tdTomato and Sox9 staining (Fig 3)** Cre recombination was induced by tamoxifen injection in 3-month-old mice. Mice were sacrificed, and one hemisphere was fixed in 4% paraformaldehyde for 1 d, stored in sucrose (15% and 25%), and embedded in Tissue-Tek (Fisher Scientific #12678646). Saggital sections (30 μm) were obtained using a cryostat (Thermo Fisher) and mounted onto slides. Brain sections were blocked with 10% normal goat serum (Vector Labs) and 0.3% Triton X-100 (Sigma) in PBS for 1 h. Subsequently, brain sections were incubated with chicken anti-GFP (1:500; Abcam, RRID: AB_300798) and rabbit anti-SOX9, 1:2000 (Millipore, RRID: AB_2239761) in 5% normal goat serum and 0.05% Triton X-100 (Roth #3051.3) in PBS overnight at 4°C followed by 3x rinsing with 5% normal goat serum (Biozol #VEC-S-1000) in PBS for 5 min. Sections were incubated with secondary antibodies from goat (anti-chicken Alexa Fluor 488 and anti-rabbit Alexa Fluor 647, 1:1,000 (Thermo Fisher) in PBS and 0.05% Triton X-100 for 3 h at room temperature, rinsed, and mounted in Fluoromount-G (Southern Biotech). Images were acquired using either a confocal laser-scanning microscope (LSM 700; Zeiss) with a 20x (NA 0.8) objective and a slide scanner (Axio Scan.Z1; Zeiss) with a 10x objective (NA 0.45), with the following filter settings: LSM700, 490–555 BP, 640 LP, 560 LP; AxioScan.Z1, 470/40 BP, 525–50 BP, 587/25 BP, 647/70 BP, 640/30 BP, 690/50 BP. The same image acquisition settings were used for each staining. Immunohistochemical data were quantified using a customized pipeline that includes the automated identification of the Sox9 positive cell nuclei and GCaMP positive cell cytoplasm using CellProfiler. **Cell type marker stainings (Fig 4)** Brain hemispheres were emersion fixed in 10% buffered formalin for two days. After cryoprotection in 30% sucrose in PBS, 40 µm coronal cryosections were taken and sections were stored until use in 50% 0.1 M PBS, 30% ethylene glycol, and 20% glycerol at −20C. Sections at the level of AP-1.46 (Franklin and Paxinos, 2013) were selected, and double incubated overnight at 4°C in primary antibodies against HA, 1:200 (clone: 3F10, Roche, AB_2314622) and the respective marker antibody. Primary antibodies (with dilutions): OSP, 1:200 (Abcam, RRID: AB_2276205), NeuN, 1:500 (Millipore, RRID: AB_2298772), Sox9, 1:2000 (Chemicon, RRID: AB_2239761), GFAP, 1:500 (Dako, RRID: AB_10013382), Iba1, 1:100 (Wako, RRID:AB_839504). Secondary antibodies (dilutions 1:1000): anti-rat IgG to detect 3F10 (Jackson ImmunoResearch 712-165-153 or 712-545-153) and appropriate second antibody (mouse: 715-545-151; rabbit: 711-225-152 or 711-585-152). Micrographs of the CA1-region of the hippocampus and deep layers of cortex immediately above CA1 were taken with a confocal microscope (LSM 700, Zeiss).

### RNA in situ hybridization

FFPE sections were prepared as for ribosomal staining sections. Prior to dewaxing sections were incubated in an oven at 50C for 1 hour. Afterwards we followed the ACDbio protocol for RNAscope 2.5 HD Assay Red and the following probes: Aldh1l1: 405891, Gja1: 486191, Slc1a3: 430781.

### Immunoblotting

RiboTag supernatants were mixed with 4x LDS (lithium dodecyl sulfate) sample buffer containing 40 mM DTT and denatured at 70°C for 10 min prior to loading on 10% NuPAGE Novex midi gels (Thermo Fisher). Gels were run using MES [2-(N-morpholino)ethane sulfonic acid] running buffer at 150-160 V (110 V for the first 15 min) and then were electro-transferred to nitrocellulose membrane (Bio-Rad), submerged in transfer buffer (20% methanol, 25 mM Tris-Cl, 0.19 M glycine), using a Criterion transfer tank (BioRad), at 0.7 A, for 70 min. Membranes were blocked for 20-30 min at room temperature (RT) in 5% powdered milk in PBS-T (PBS with 0.05% Tween-20) and then incubated with primary antibody diluted in blocking buffer overnight at 4°C. Next, blots were washed 4x with PBS-T and incubated with secondary antibody for 30-60 min at RT, followed by 5x PBS-T washes and imaging with the Li-Cor Odyssey imaging system (Li-Cor). For re-probing, membranes were stripped for 10 min with Re-Blot strong solution (EMD-Millipore), followed by extensive washing and re-blocking. Primary antibody: anti-Slc1a3 (dil. 1:2000, CST, clone D20D5, RRID: AB_10694915), rabbit-anti-β-actin (dil. 1:10.000, Sigma, RRID: AB_476697). Secondary antibodies: donkey-anti-rabbit IRDye 680RW (dil. 1:10.000, Li-Cor, RRID: AB_10706167), donkey-anti-mouse IRDye 800CW (dil. 1:20.000, Li-Cor, RRID: AB_1850023).

### Translatome purification with RiboTag

We previously optimized our RiboTag protocol to increase its specificity (Kaczmarczyk et al., 2019). For RiboTag experiments mice were sacrificed at 50 days of age. Brain was removed, and olfactory bulb was cut out, and then cerebellum separated from cerebrum. deep frozen brain tissue was used to prepare homogenate in Polysome Buffer (PSB) containing 50 mM Tris (pH 7.5), 100 mM KCl, 12 mM MgCl2 and 1% Nonidet P-40, 1 mM DTT, 100 U/ml Ribolock RNase inhibitor, 100 µg/ml cyclohexamide and 1 tab/5 ml SigmaFast protease inhibitor cocktail with EDTA (Sigma). One hemisphere was used per replicate. Homogenates were prepared with 1 ml and 2 ml of PSB was used per cerebellum (both hemispheres were pooled) and cerebrum (the remaining part of the brain left after removal of cerebellum and olfactory bulb, one hemisphere) respectively, in Potter-Elvehjem homogenizers using a motorized (∼450 rpm) pestle and then cleared by centrifugation (10.000 g / 10 min / 4°C) to remove nuclei and cell debris to obtain the supernatant (S1). 50 μl of lysate was immediately mixed with Qiazol and purified as input. Supernatant was pre-cleared with Protein G Dynabeads (PGDB, Thermo Fisher) for 30 min at 4°C on a rotator and S1 (1 ml for cerebellum and 1 ml for cerebrum) was incubated with 5 μg of anti-HA mAb (Roche, clone 12CA5, RRID: AB_514505) at 4°C for 60 min on a rotator. Sample was then transferred to prepared equivalent of 37.5 μl total bead suspension PGDB (PSB-equilibrated) and incubated as above for additional 100 min. Afterwards, both Ribo-Tag and Ago-Tag beads were washed 3 × 5 min in High Salt Buffer (HSB) containing 50 mM Tris (pH 7.5), 300 mM KCl, 12 mM MgCl2 and 1% Nonidet P-40, 1 mM DTT, 50 U/ml Ribolock RNase inhibitor (Thermo Fisher), 100 µg/ml cycloheximide, 1 tab/20 ml of SigmaFast protease inhibitor cocktail with EDTA (Sigma) and additional 3 × 5 min in Extra High Salt Buffer (EHSB, identical to HSB but containing additional 300 mM NaCl). During each wash, beads were rotated gently at 4°C. Following removal of the last wash solution, 700 μl Qiazol (Qiagen) was added and the beads were incubated for 15 min at RT with vigorous (>1000 rpm) agitation. RNA was extracted from input and RiboTag samples using miRNeasy Micro kit (Qiagen) and eluted with 28 μl of water.

## Acknowledgements

This work was supported by generous internal funding from the Wallenberg Center for Molecular Medicine and the German Center for Neurodegenerative Diseases. We thank Alessandro Petese for excellent technical assistance and Wieslaw Krzyzak and Joachim Degen for performing blastocysts injection.

