## Supplementary data for "Insights into calcium signaling and gene expression in astrocytes uncovered with 129S4 Slc1a3-2A-CreERT2 knock-in mice"

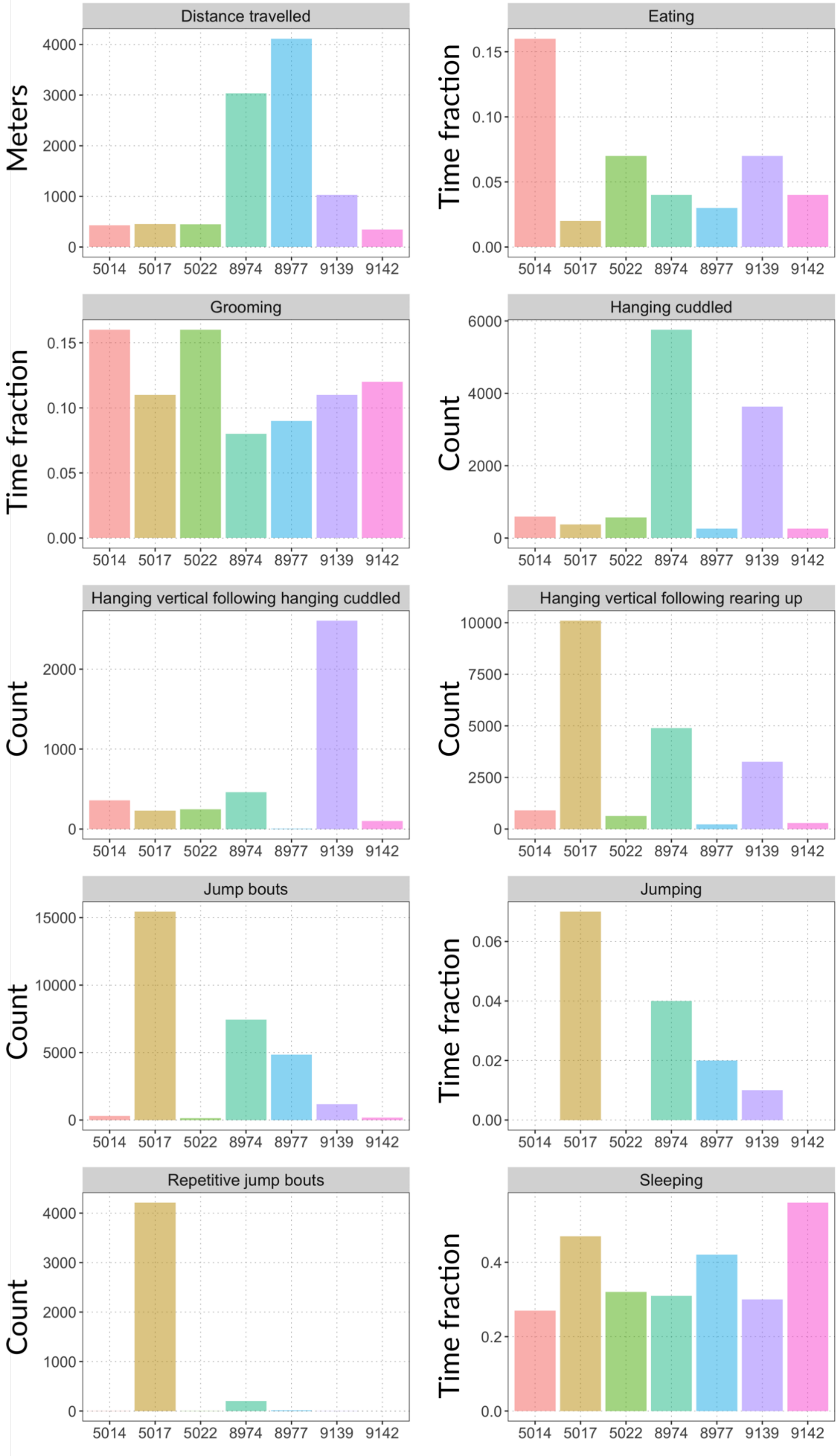


**Figure S1. Comparison of individual C57Bl/6 mice from Figure 1A across behavioral categories.** Top-ranked mice (color-coded) in each behavioral category were cross-compared across the categories. Numbers on x-axis are mice IDs. Total number of mice is smaller than the number of categories because typically the same mouse was top-ranked for the related categories (e.g., hanging-related, jumping-related, etc.,).

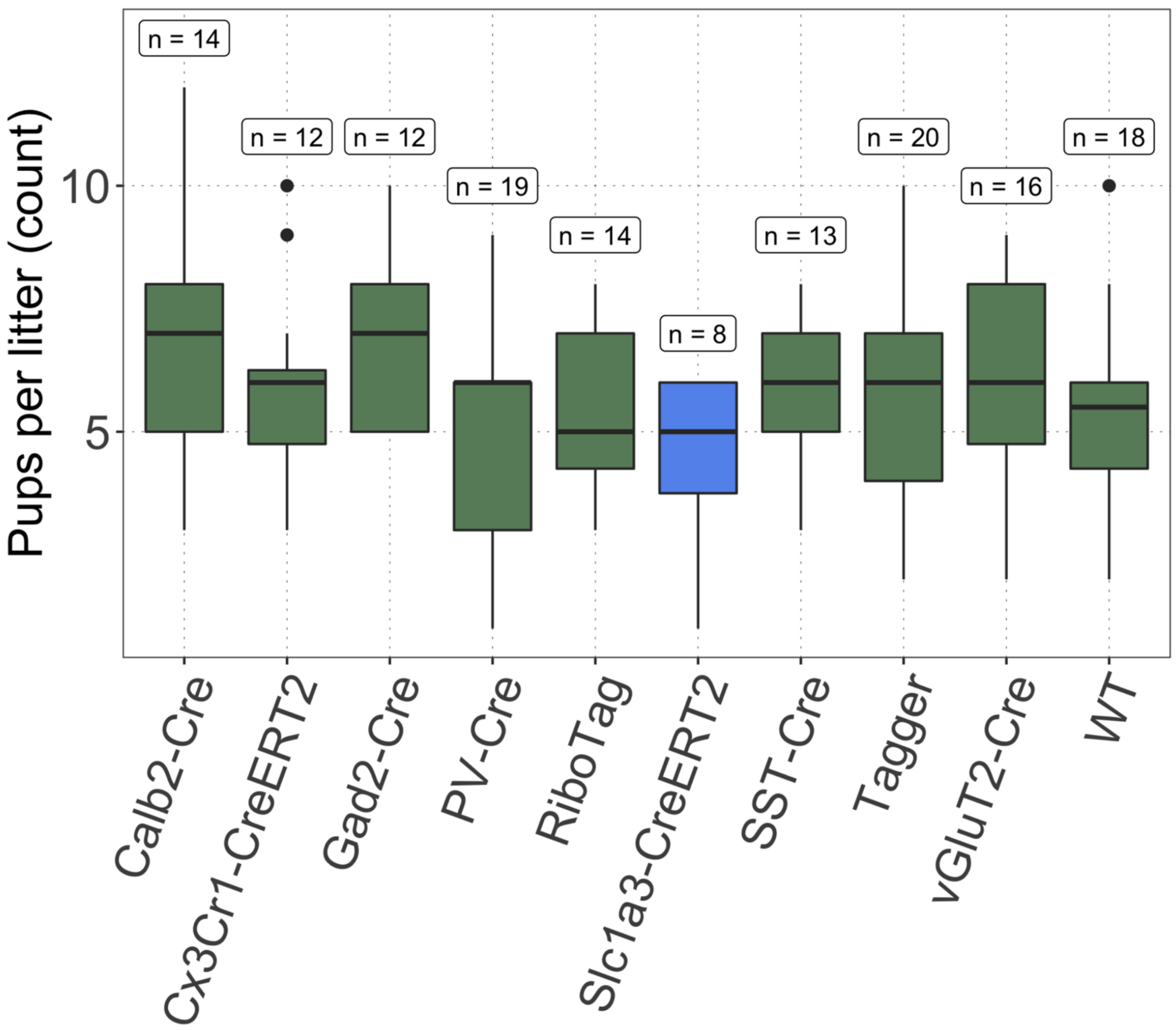


**Figure S2. Litter sizes of S4 lines.** The breeding productivity of ten S4 lines in our colony (displayed as the number of pups per litter) produced over a period of 1.5 years. Differences were not significant (one-way ANOVA). By comparison, B6 breeding outputs in the same animal facility in three independent colonies with multiple lines resulted in an average of 4.7 pups from 61 productive matings (15 were unproductive). n denotes the number of animals included in the statistic.

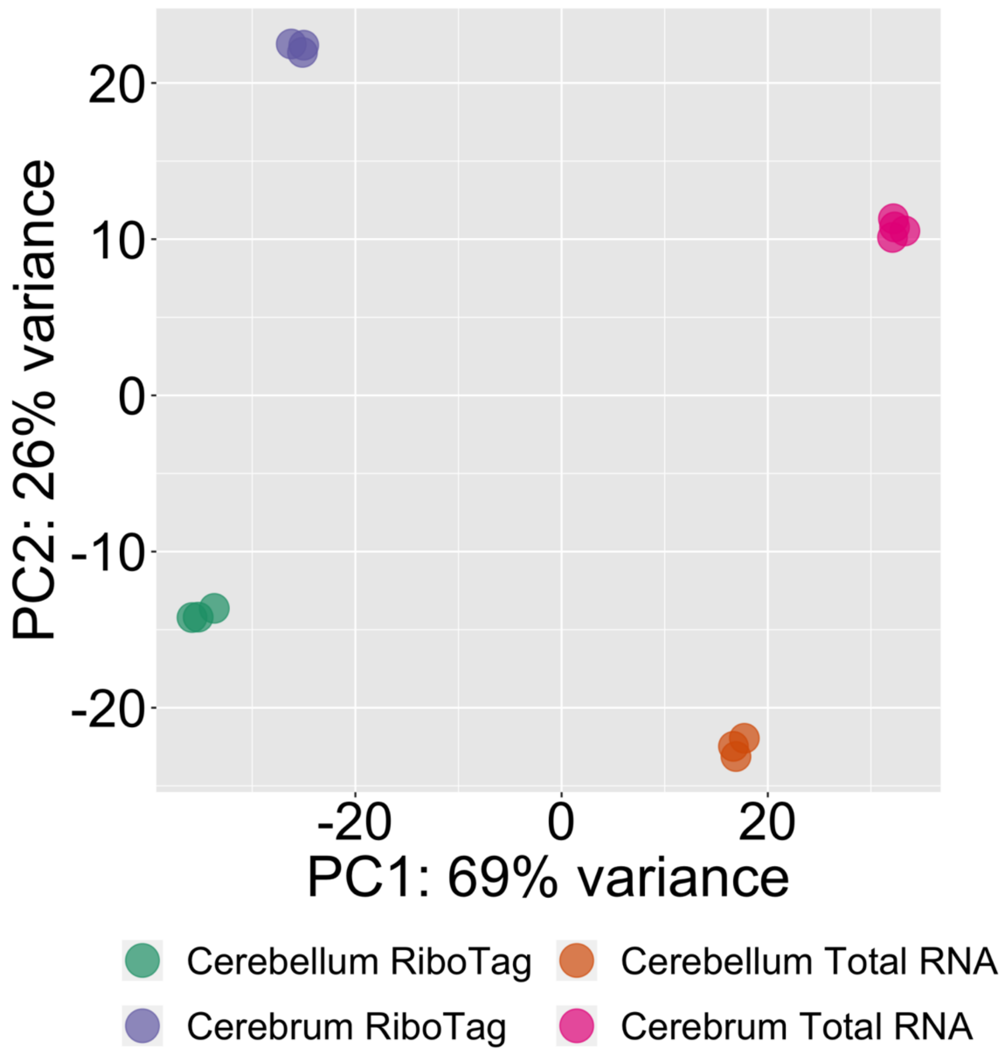


**Figure S3.** Principal component analysis of the RiboTag IP and input RNA data showing that the RiboTag purifications were reproducible.

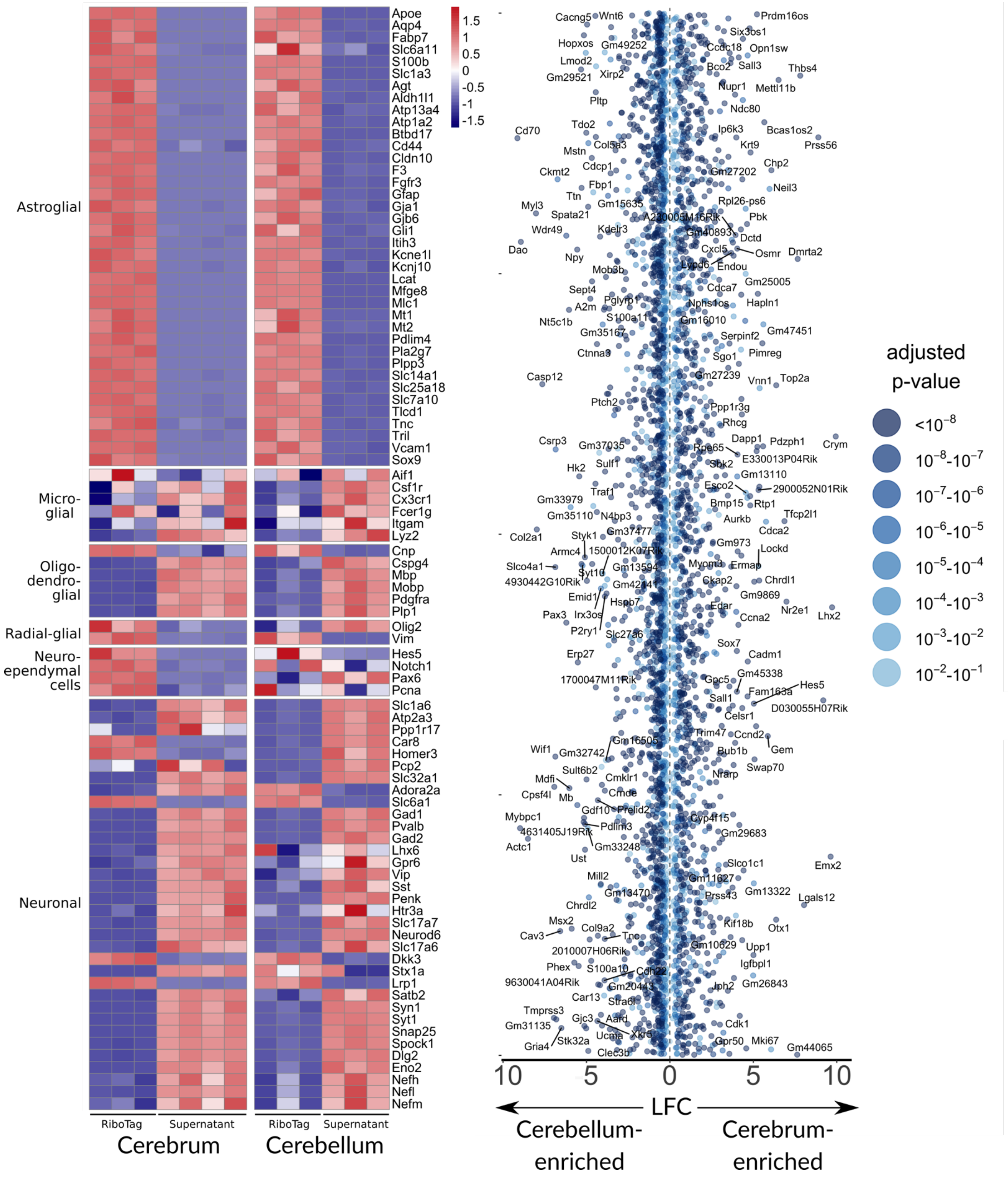


**Figure S4.** (**A**) **Heatmap of extended list of marker genes.** Each column represents single biological replicate. Z-score was calculated as follows: $Z = (x - \text{meanrow}(x))/SD(\text{row})$, where SD is standard deviation. (**B**) Dot plot with significantly different (FDR < 0.05) genes between BG and AC. Vertical dimension is random scatter. 100 top genes in BG and 100 top genes in AC (by LFC) were labeled. Color scale reflects FDR.

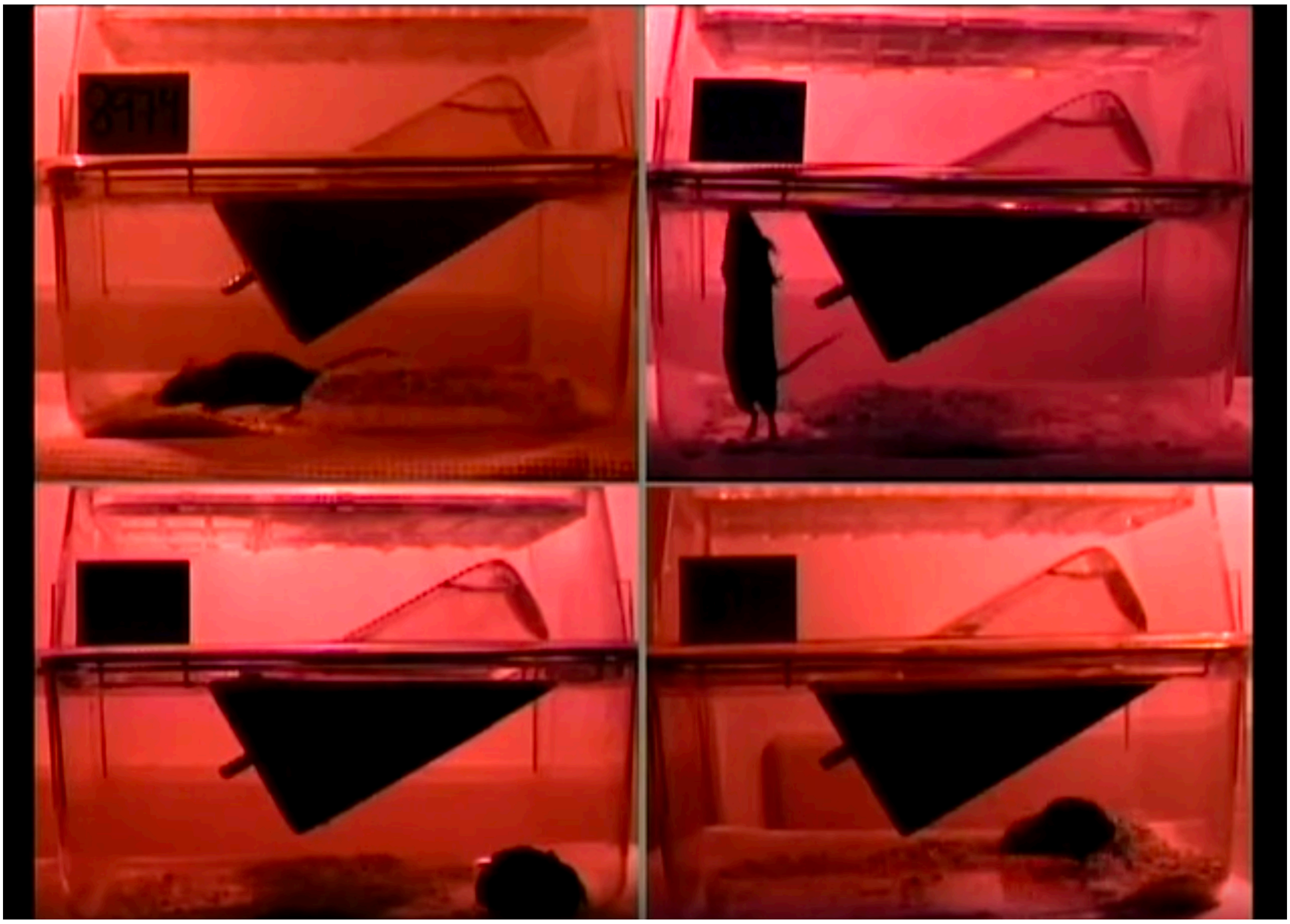

**Movie S1.** Representative littermate C57Bl/6 mice observed at night (https://tinyurl.com/wcmz9zf).

| ID (as on Fig B2) | Name | Sequence | Binding to | Orientation |
|---|---|---|---|---|
| P1 | Cre_cDNA_for | GCATTACCGGTCGATGCAACGAGTG | CreERT2 | F |
| P2 | Cre_cDNA_rev | GAACGCTAGAGCCTGTTTTGCACGTTC | CreERT2 | R |
| P3 | Slc1a3_genomic_for | CAGCTCCTCCTGTATCCAGTGTTCT | Intron 9 | F |
| P4 | Slc1a3_genomic_rev | TACTCCCCCGCAGCCTAGTGTTA | Exon 10 | R |
| P5 | Slc1a3_rev | GAACAGTTTCCAACACTTGGTGCT | Exon 10 | R |
| P6 | Cre_for | GACACTTTGATCCACCTGATGGC | CreERT2 | F |
| P7 | Slc1a3_for | GGTCCTCACTGTTCACCTTCTGT | Intron 9 | F |

**Table S1.** PCR oligos for genotyping and verification of gene targeting.
